# A Functional Analysis of a Resorbable Citrate-based Composite Tendon Anchor

**DOI:** 10.1101/2024.01.08.571095

**Authors:** Arun Thirumaran, Anup Poudel, Magesh Sankar, Meletios Doulgkeroglou, Jeremiah Easley, Ben Gadomski, Manus Biggs

## Abstract

Orthopaedic fixation seeks solutions to the challenges of non-union, reconstructive surgery, and soft tissue injuries by providing stability and tissue alignment during the healing process. Risks associated with fixation devices such as peri-implant resorption, implant loosening and sub-optimal device resorption remain a significant challenge in the development of transient fixation devices. Osteomimetic biomaterials, and in particular, bio-resorbable polymer composites designed to match the mineral phase content of native bone have been shown to exhibit osteoinductive and osteoconductive properties *in vivo* and have been used in bone fixation for the past 2 decades. However, the specific signalling pathways driving the osteogenic response to these biomaterials remain largely unknown.

In this study a resorbable, bioactive, and mechanically robust citrate-based composite, formulated from poly(octamethylene citrate) (POC) and hydroxyapatite (HA) (POC-HA) was investigated as a potential orthopedic biomaterial. *In vitro* analysis with human Mesenchymal Stem Cells (hMSCs) indicated that POC-HA composite materials supported cell adhesion, growth, and proliferation and increased calcium deposition, alkaline phosphatase production, the expression of osteogenic specific genes and activation of canonical pathways leading to osteoinduction and osteoconduction. Further *in vivo* evaluation of a POC-HA tendon fixation device in a sheep metaphyseal model indicated the regenerative and remodelling potential of this citrate-based composite material in orthopedic applications.

## 1. Introduction

A significant goal of orthopaedic therapies is to reduce and stabilise a fracture or defect through techniques and devices to promote healing and complete recovery of structure and function. Orthopaedic fixation, in particular, relies on mechanically robust and biocompatible screws, anchors or plate systems to enhance bone stability or promote bone/soft tissue apposition. Advances in materials science have accelerated the development of bioresorbable and bioactive fixation devices which undergo controlled degradation and regulate the critical processes of osteoinduction, osteoconduction and osseointegration.^[1]^ In particular, tendon anchoring devices have revolutionized orthopedic surgery allowing for rapid and efficient tendon fixation to a bone following surgery of the joints of the upper or lower limbs

Previous studies into the development of effective resorbable orthopaedic devises have reported on the use of bioresorbable composite materials, employing a microscale mineral phase dispersed within a polymer matrix. In particular, synthetic polymer composite formulations such as Polycaprolactone (PCL)^[2–4]^, Polymethyl Methacrylate (PMMA)^[5,6]^, Poly-lactic acid (PLA)^[7,8]^, Poly-L-lactic acid (PLLA)^[9]^, Polyglycolic acid (PGA)^[10]^, their copolymer polylactic-co-glycolic acid (PLGA)^[11,12]^, and Poly(glycerol-succinate) (PGS)^[13]^ containing tricalcium phosphate (TCP) or hydroxyapatite (HA) particulates have been extensively described and have seen clinical success as scaffolds for orthopaedic regenerative engineering applications^[10,14]^. Furthermore, recent studies indicate that the chemistry of these materials may be tailored to improve the biocompatibility, mechanical, osteoinductive and osteoconductive properties of these composite materials *in vitro* and *in vivo*^[15–18]^.

However, there are recognised shortcomings with the use of synthetic composites as orthopaedic materials, specifically related to their manufacturing processes, bioactivity, immune response, biosorption, and osteoinductive capacity. Addressing the limitations of current composites materials used in the manufacture of orthopaedic fixation devices, citrate-based elastomeric composites ^[19]^ have been identified as alternative resorbable and biocompatible ^[20]^ ^[21–23]^ alternative materials with mechanical properties comparable to that of human cortical bone^[19]^. Citrate, a critical intermediate in the Krebs cycle, is highly concentrated in native bone (90% of body’s total citrate content is in the skeletal system) and is reported to be closely associated with bone metabolism and formation. Citrate not only serves as a calcium-solubilizing agent^[22,24]^, but also plays an essential role in the anatomy and physiology of bone apatite nanocrystals^[24]^, which regulate and provide bone with unique load-bearing properties^[25]^. Critically, citrate-based composite materials formulated with HA microparticles are extensively reported to promote osteogenic differentiation of MSCs *in vitro* ^[26]^ and to promote osteochondral repair processes in pre-clinical animal models[23,27–29].

In this study a citrate-based composite, poly(octamethylene citrate) (POC) was synthesized using two acid to diol ratios (POC 1:1.1 and POC 1:1.3) and formulated into a resorbable composite material (POC-HA) through the addition of hydroxyapatite. The mechanical, degradation and biocompatibility properties of the POC-HA citrate composite materials were assessed relative to a control (PLDLA-TCP) biodegradable composite, commonly used in FDA approved biodegradable orthopaedic devices. Furthermore, the potential of these materials in activating hMSC osteo-responsive signaling pathways was assessed *in vitro* via RNA sequencing and computational pathway analysis. Finally, POC-HA composites were fabricated into suture anchors, implanted into a sheep metaphyseal defect model and assessed in vivo.

POC-HA composites were observed to enable cell adhesion, proliferation and differentiation of hMSCs into osteogenic lineage as evident by the deposition of calcified extracellular matrix (ECM), production of alkaline phosphatase (ALP) and expression of osteogenic marker osteopontin (OPN). In addition, hMSCs cultured on POC-HA composites underwent an increase in osteoresponsive signaling pathway activation leading to osteoinduction, osteoconduction, and ECM deposition, maturation and mineralisation. *In vivo* pre-clinical evaluation of POC-HA composites in a sheep metaphyseal model further supported the osteogenic capacity of the POC-HA composite material *in vivo* and enhanced peri-implant integration was observed relative to a control PLDLA-TCP anchor. Together, this study presents a comprehensive in vitro and in vivo analysis of the functional response to a citrate-derived composite tendon anchor.

### Mechanical analysis of POC-HA and control PLDLA-TCP composites

In order to assess the compressive mechanical properties of the formulated citrate-based composites, uniaxial compressive analysis of POC 1.1-HA, POC 1.3-HA and control PLDLA-TCP composites was performed under ambient conditions using a strain rate of 1.3 mm/min. POC 1.1-HA and POC 1.3-HA composites demonstrated a similar stress-strain responses (fig. 1a) typical of an elastomeric material exceeding 40% strain at peak stress. In contrast, the stress-strain responses of the PLDLA-TCP composite showed plastic deformation and underwent less than 10% strain at peak stress. The compressive modulus (fig. 1b) of PLDLA-TCP composites was significantly higher than POC-HA composites with no significant difference between POC 1.1-HA and POC 1.3-HA composites observed. Conversely, the peak stress of both citrate-based composites was significantly higher than PLDLA-TCP. Interestingly, POC 1.1-HA composites showed a higher peak stress and strain (fig. 1c&d) than POC 1.3-HA possibly due to an increase in the available carboxylate chemistry for polymer-bioceramic interactions.

**Figure 1:**
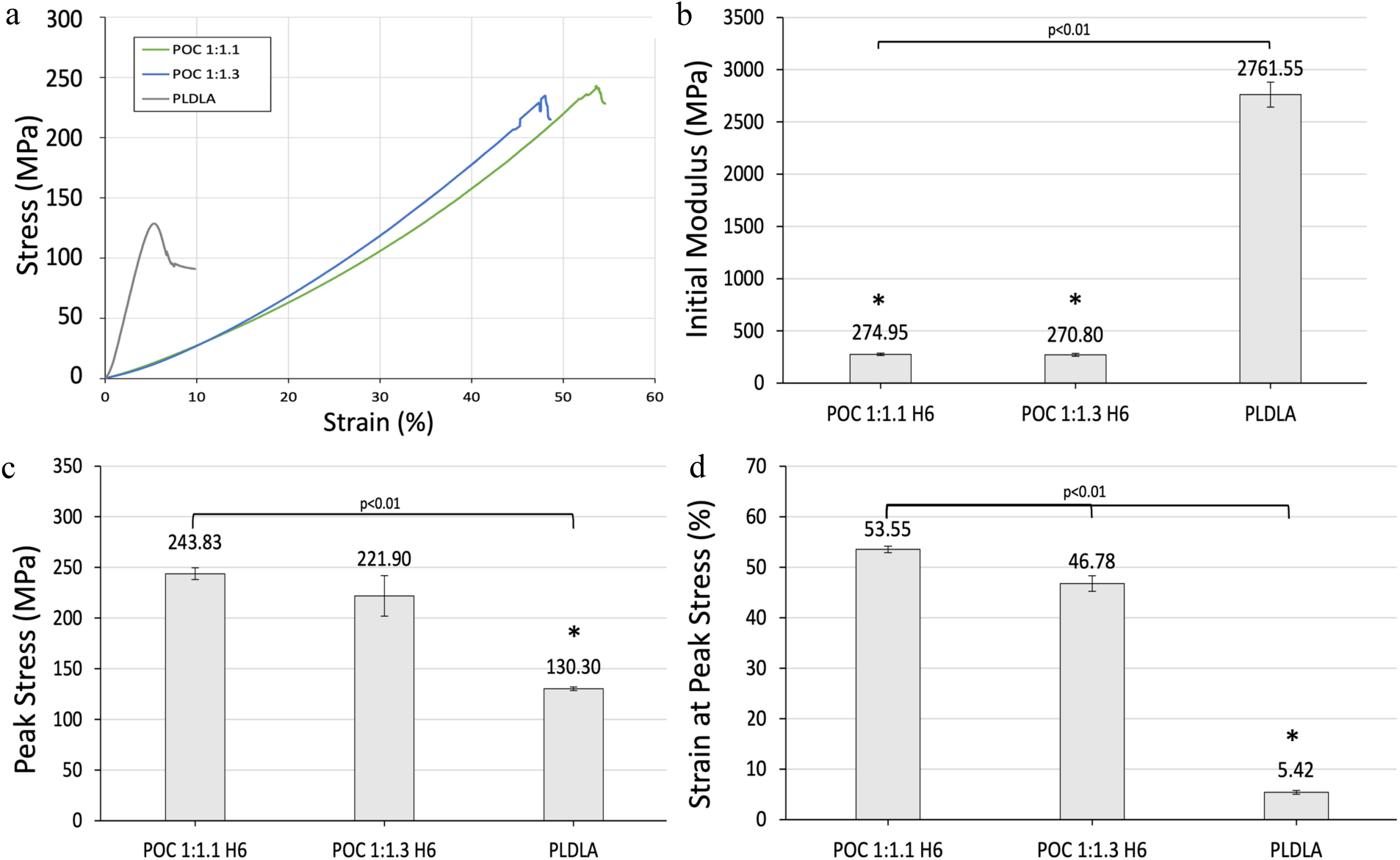
Uniaxial compression testing of POC-HA and PLDLA-TCP composites A) representative stress-strain curves, B) initial modulus, C) peak stress, and D) strain at peak stress. Data is represented as mean ± standard deviation. n=4, * indicates p value of significance <0.01.

### Degradation analysis of POC-HA and control PLDLA-TCP composites

White light interferiometry was performed to assess the surface topography of POC 1.1-HA, POC 1.3-HA, and PLDLA-TCP composite materials before and after cell seeding.

Non-degraded (day 0) PLDLA-TCP composites possessed a heterogenous surface morphology with relatively large pit and peak features (fig. 2a) resulting in an increased roughness relative to both POC formulations at day 0 (fig. 2b&c). PLDLA composites were observed to undergo surface degradation 7 days post MSC seeding (fig. 2d), as identified through a significant increase in surface feature peak-pit depth and roughness (fig. 2g). Conversely The roughness of 1.1 and 1.3 POC formulations were not observed to significantly change by day 7 (fig. 2e&f, 2g).

**Figure 2:**
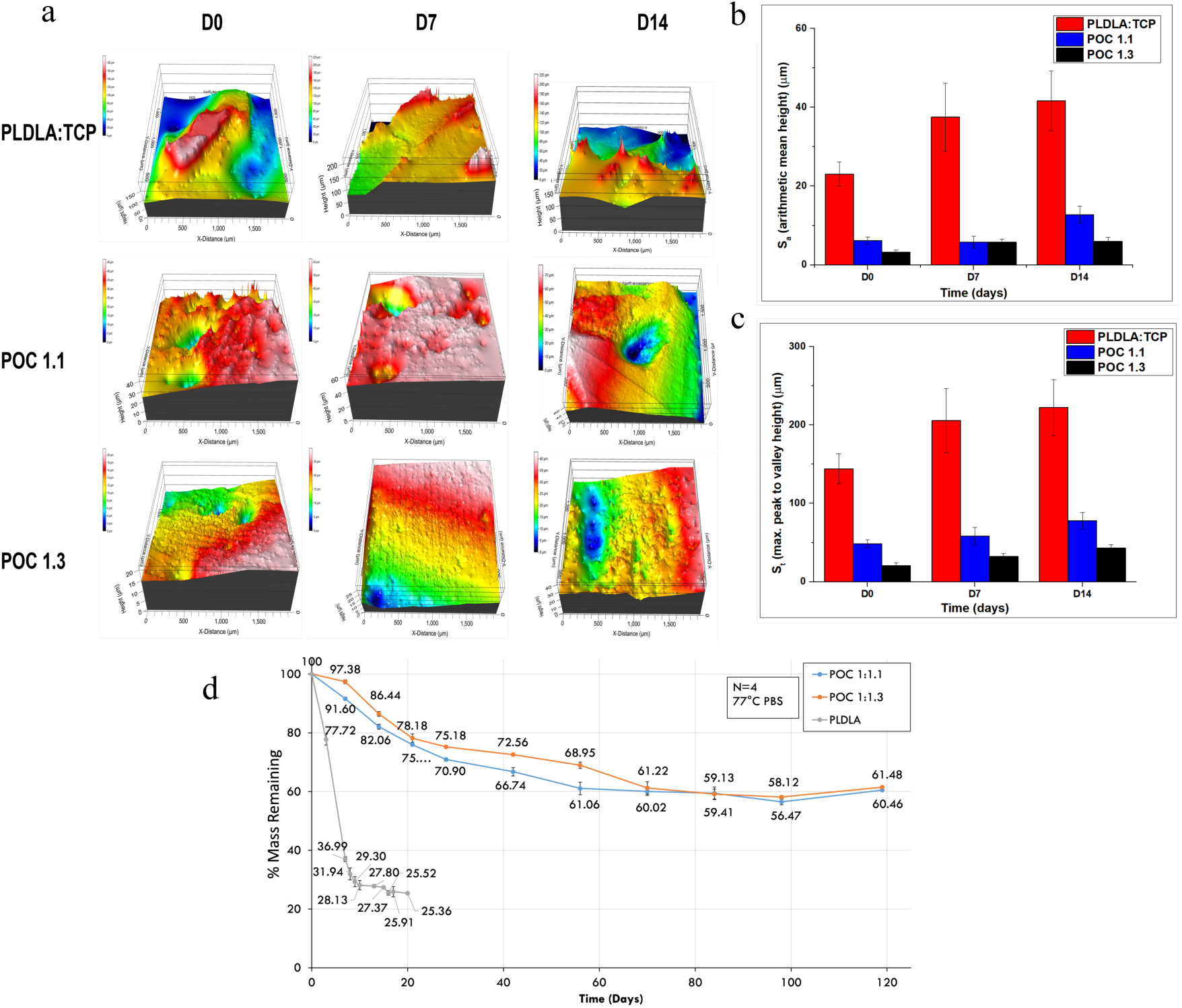
The effects of in vitro degradation on the surfaces roughness and mass loss of POC 1.1, POC 1.3 and PLDLATCP substrates in vitro. Surface profilometrey imaging (a) and analysis (b and c) of roughness (S_t_, S^a^,) of PLDLATCP, POC 1.1 and POC 1.3 substrates for pristine and D7 (PBS) substrates immersed in PBS for up to 14 days at 37°C. Accelerated degradation analysis of mass loss of POC 1.1, POC 1.3 and PLDLA-TCP substrates immersed in PBS at 77°C for up to 120 days.

Polymer hydrolysis was conducted under accelerated conditions at 77°C in phosphate buffered saline (PBS) for POC-HA and PLDLA-TCP composites. PLDLA-TCP composites displayed rapid polymer degradation in these conditions with complete polymer loss after 20 days. In contrast, POC composites displayed complete polymer hydrolysis after 70 days (fig. 2h). POC 1.3 composites initially resisted polymer hydrolysis when compared to 1.1 composites due to the enhanced polymer chain crosslinking and reduction in hydrophilic carboxylate chemistry.

### 1.13 Differential scanning Calorimetry (DSC) of POC-HA and PLDLA-TCP composites

Figure 3 shows the DSC heat flow curves for POC 1.1-HA, POC 1.3-HA and control PLDLA-TCP composites at 0, 7 and 14 days post hMSC seeding. The onset of glass transition for POC 1.1-HA materials (fig. 3a) was observed to occur at -41°C in non-degraded samples, was shifted to ca.-21°C at 7 days post degradation, and was not present by 14 days post cell seeding. The sizeable leathery region (-41.94 to -0.91°C) of the polymer suggested a wide variation of chain molecular weight within the polymer matrix. The shift in the onset of glass transition temperature to -21°C at 7 days post cell seeding and the absence of this transition temperature at 14 days post cell seeding suggested a degradation of the polymer as a function of molecular weight. A notable endothermic peak at 110°C progressing above the baseline at ca. 150°C was observed suggesting thermal degradation.

**Figure 3:**
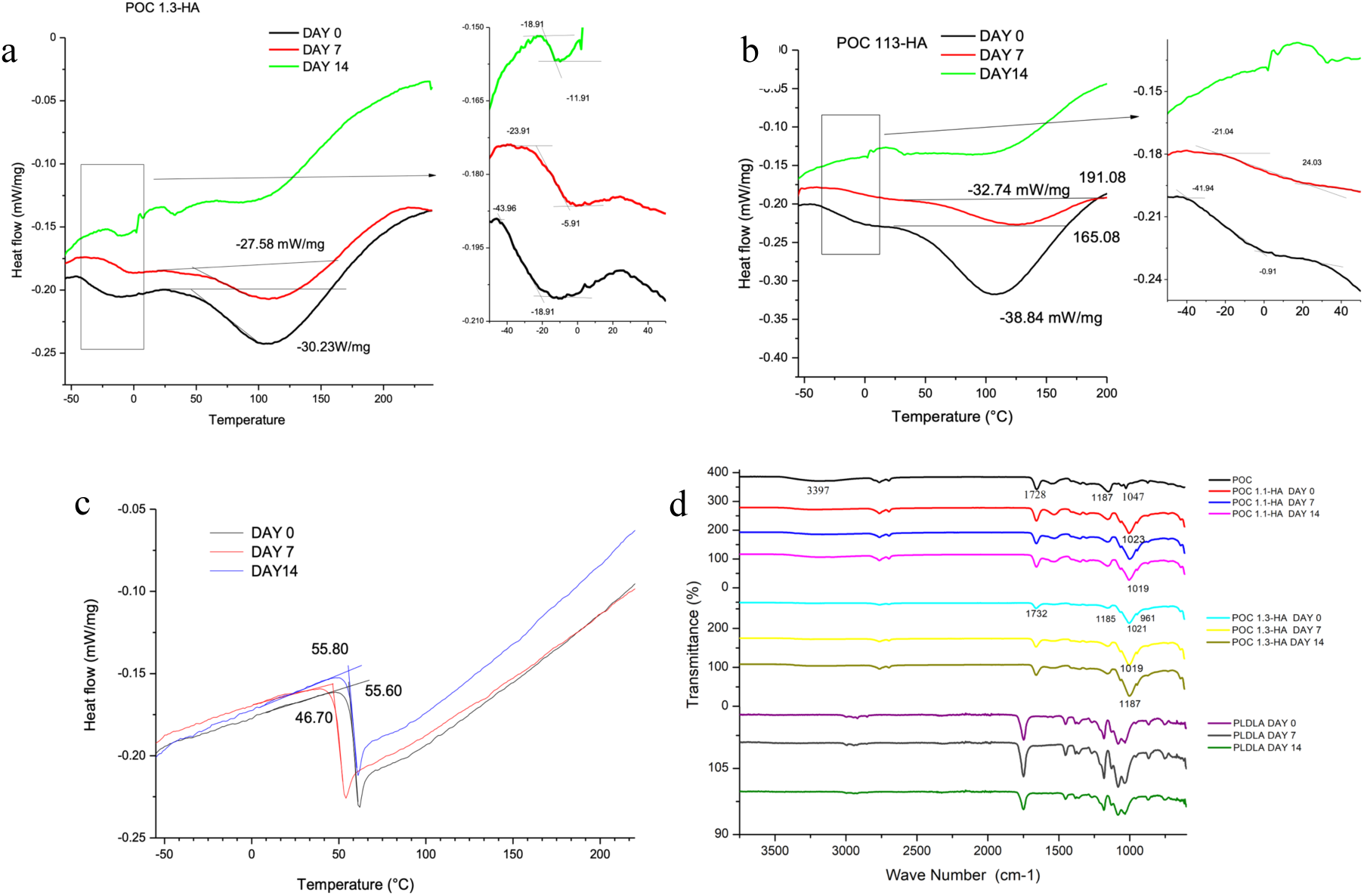
DSC analysis of (a) POC 1.1-HA, (b)POC 1.3-HA, and (c) PLDLA-TCP, 0, 7 and 14 days post cell seeding. FTIR Spectra of POC 1.1-HA POC1.3-HA and PLDLA-TCP pre-cell seeding (day 0), and on day 7 and day 14 post-cell seeding (d).

POC 1.3-HA composites also demonstrated a similar trend (fig. 3b), however, this composite demonstrated increased stability relative to POC 1.1-HA and a glass transition phase was still present at 14 days post cell seeding. There were no significant changes in the onset of the glass transition temperature or the glass transition region as a function of cell culture duration between the POC 1.1-HA and POC 1.3-HA composites. Conversely, PLDLA-TCP composites demonstrated amorphous behaviour and a glass transition temperature of 55°C (fig. 3c) which was observed to decrease by day 7, followed by an increase at day 14, suggesting a non-linear degradation process.

### 1.14. Fourier Transform Infrared (FTIR) Spectroscopy of POC-HA and control PLDLA-TCP composites

FTIR analysis of POC 1.1-HA and POC 1.3-HA composites was performed on pristine samples (day 0), and at 7 and 14 days following hMSC seeding to assess cell-mediated degradation of nanocomposite surface chemistry (fig. 3d). FTIR spectra were characteristics for POC elastomer chemistries and bands were observed at 1728 cm^-1^ (C==O stretching), 1615 cm^-1^ (strong C=C carbonyl stretching), 1393 cm^-1^ (symmetric stretches of β-COO−), 1220 cm^-1^ (C-O stretching), 1079 cm^-1^ (stretching vibration of C-O-C bond), 1047 (absorption peaks of NO), 880 cm^-1^ and 722 cm^-1^ (C=C bending). The broad peaks centred at 3397 cm^-1^ was assigned as hydrogen-bonded hydroxyl groups. Similarly, the peak at 2930 cm^-1^ was assigned to the methylene groups of the citrate-based elastomer.

The addition of HA to POC polymer shifted the 1728 cm^-1^ peak to 1732 cm^-1^, the 1615 cm^-1^ peak to 1595 cm^-1^, the 1393 peak to 1398 cm^-1^, the 1220 cm^-1^ peak to 1187 cm^-1^, the 1079 cm^-1^ peak to 1023 cm^-1^ and changed the intensity of multiple peaks suggesting the interaction of HA with C==O, C=C, COOH, C-N, C-O and C-O-C. The shift of the peak at 1023 cm^-1^ to 1019 cm^-1^ assigned to stretching vibration of C-O-C bond in both citrate-based composites at later time points, suggests degradation of the polymer molecule. Fig. 3d also shows the FTIR spectra of PLDLA-TCP at day 0, 7 and 14 following hMSC seeding. The characteristic polylactide bands of PLDLA were observed at 1033, 1081 and 1267 cm–1 (=C-O stretching), 1360 and 1381 cm–1 (CH2 wagging), 1451 cm–1 (CH3 bending), 1747 cm–1 (C=O stretching in the ester group), 2918 cm–1 (CH stretching) and 2994 cm–1 (CH3 stretching). Shifts in polylactide bands at 1033, 1081 and 1267 (=C-O stretching), 1360 and 1381 cm–1 (CH2 wagging), 1451 cm–1 (CH3 bending), 1747 cm–1 (C=O stretching in the ester group) were observed. Previous research suggests these minor shift are due to a change in microphase domains due to copolymer-solvent interaction. This behaviour is also supported by DSC analysis. The shift in wavelength observed at 7^th^ day in PLDLA-TCP spectra was again observed at day 14 post hMSC seeding.

### 1.15 In vitro Analysis of POC-HA Cytocompatibility

Calcein staining of hMSCs (Figure 4 a-d) indicated that pristine POC formulations without bioceramic addition did not support cell adhesion and growth while POC 1.1-HA, POC 1.3-HA, and PLDLA-TCP composites supported cell adhesion and proliferation for up 21 days in vitro. Alamar blue assay indicated that hMSCs cultured on the pristine POC formulations demonstrated a significant decrease in metabolic activity at all time points (Figure 4e), while there was no significant difference in the metabolic activity of hMSCs culture on both POC 1.1-HA and POC 1.3-HA composite formulations at any experimental time points. Conversely, hMSCs seeded on the control PLDLA-TCP composite showed a statistically significant increase in metabolic activity at 7 and 14 days relative to both POC-HA formulations. This trend continued until day 28, with respect to the POC 1.1-HA composite (Figure 4e).

**Figure 4:**
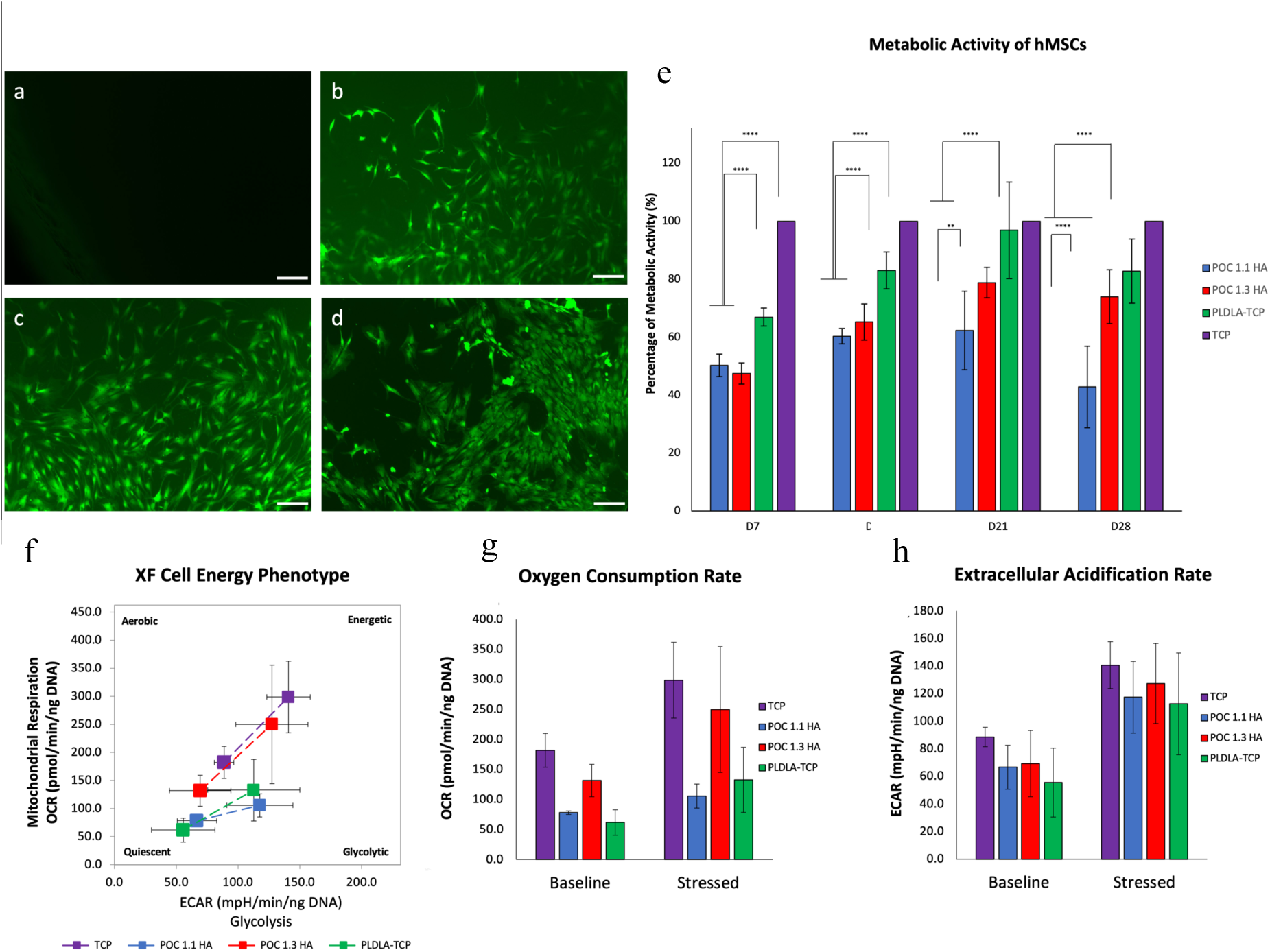
Representative images of Calcein staining showing adhesion and growth of hMSCs on POC (a), POC 1.1-HA, (b), POC 1.3-HA (c) and PLDA (d). Scale bar 200μm. Metabolic activity ofh MSCs cultured on POC-HAs, PLDLA and tissue culture plastic for 7, 14 and 21 days normalised to cells in tissue culture plastic (e) hMSCs cultured on POC 1:1.3 composites demonstrated increased cell energy phenotype relative to POC 1:1.1 and PLDLA-TCP control substrates (f). This trend was reproduced with respect to oxygen consumption rate (g). No significant differences in extracellular acidification rate were observed between any of the experimental or control groups (h). n=3, p<0.05

Further metabolic analysis was performed using an Agilent Seahorse XFP cell analyzer to assess the energy status resulting from mitochondrial respiration and glycolysis of hMSCs exposed to POC-HA and PLDLA-TCP composite conditioned media (Figure 4f). POC-HA composites induced an increase in cell energy phenotype when compared to PLDLA-TCP composites. hMSCs exposed to osteospecific media demonstrated the highest energy status (baseline OCR 182.15 OCR pmol/min/ng DNA, baseline ECAR 88.69 mpH/min/ng DNA, CR 398.94 pmol/min/ng, stressed ECAR 140.78 mpH/min/ng DNA) relative to hMSCs exposed to POC-HA and PLDLA-TCP composite conditioned media (Figure 4f). Two further parameters of cellular energy status, oxygen consumption rate (OCR) (Figure 4g) and extracellular acidification rate (ECAR) (Figure 4h) were measured during the assay. The hMSCs oxygen consumption rate was observed to significantly decrease on POC 1:1.1 and control PLDLA substrates relative to TCP controls under baseline and stressed conditions (Figure 4g). No significant differences were observed in ECAR between any of the experimental groups (Figure 4h).

### 1.16 In vitro Assessment of POC-HA Osteogenic Potential

The osteogenic differentiation of hMSCs cultured on POC-HA and PLDLA-TCP composites was assessed through the expression of osteogenic marker osteopontin, the quantification of ALP production and the deposition of mineral calcium under normal and osteogenic culture conditions.

Quantification of gene expression revealed increased OPN production in MSCs cultured on both POC-HA composites relative to hMSCs cultured on PLDLA-TCP control composites under both normal and osteogenic conditions and at all investigated time points (Figure 5A). Interestingly, by day 14, OPN expression of MSCs cultured on POC-HA composites under normal growth medica conditions was comparable to OPN expression of MSCs cultured on control TCP conditions under osteogenic growth conditions (Figure 5a).

**Figure 5:**
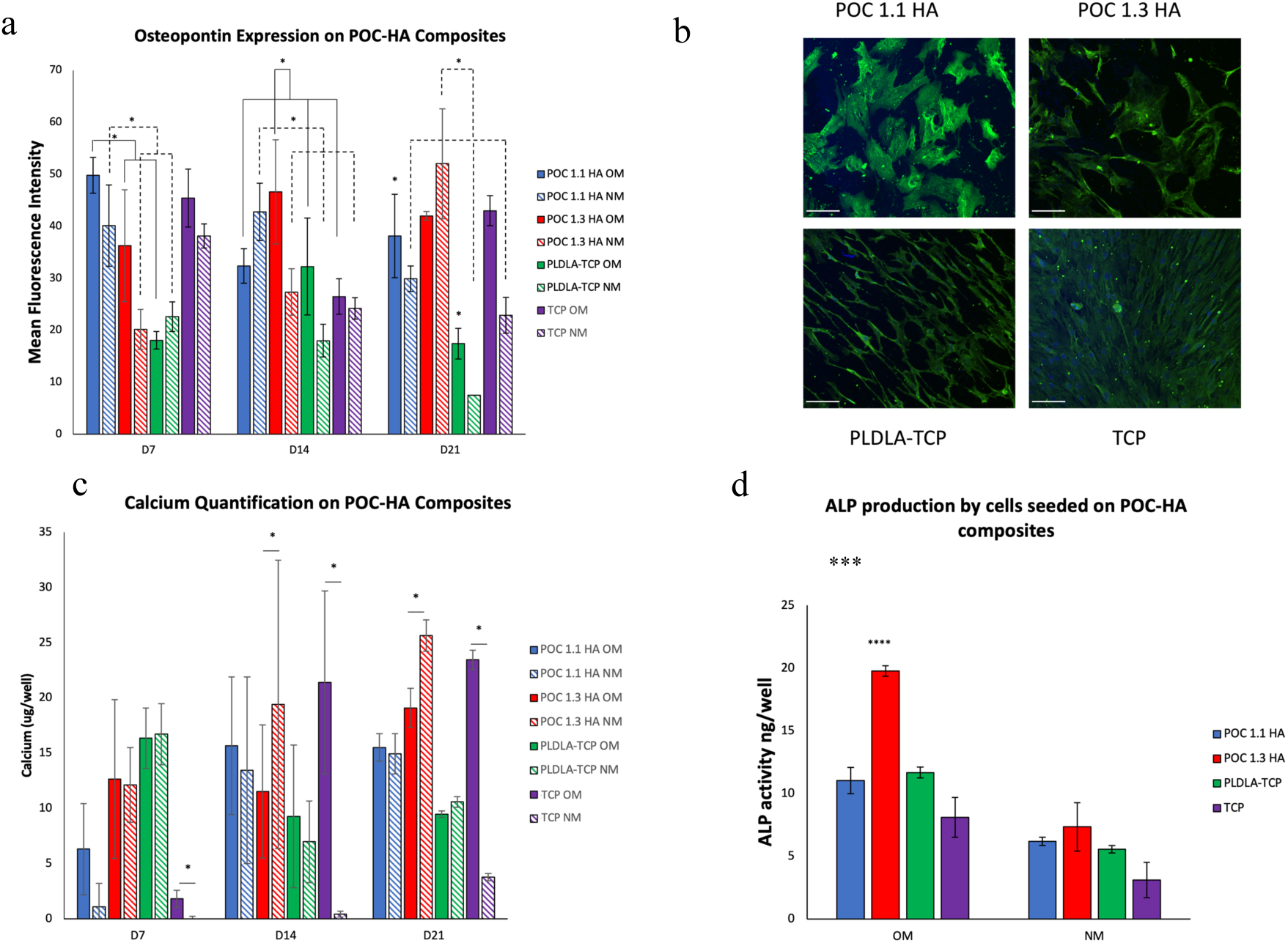
The in vitro osteospecific response of hMSCs cultured on POC-HA composites. Quantification of osteopontin expression (a) and immunostaining of osteopontin expression (b) in hMSC cultured on POC-HA and PLDLA-TCP composites in normal growth medium (NM) at day 21. Green osteocalcin, blue nucleus. (Magnification 20x, sale bar 100μm. Calcium deposition (c) of hMSCs cultured on POC-HA and PLDLA-TCP composites in either osteogenic induction media (OM) or normal growth medium (NM) for 7, 14 and 21 days. Quantification of ALP activity measured on day 14 h in MSCs cultured in OM (d).

Quantitative analysis of calcium deposition in hMSCs cultured on POC-HA and the PLDLA-TCP composites in osteogenic induction media and normal hMSC growth media for 7, 14 and 21 days indicated a time-dependent increase in calcium deposition on both POC-HA composites relative to cells cultured on the control PLDLA-TCP composite (Figure 5b). hMSCs cultured on POC 1.1-HA and POC 1.3-HA composites under osteogenic induction and normal growth media showed increased calcium deposition relative to PLDLA-TCP control composite materials at days 14 and 21. Critically, by day 14 and 21, hMSCs cultured on POC 1.3-HA composites in normal growth medium demonstrated greater calcium deposition than cells cultured in osteogenic induction media. hMSCs cultured on control PLDLA-TCP composites, demonstrated no significant difference in calcium deposition between osteogenic and normal media conditions at any time point, but demonstrated a relative decrease in calcium deposition over time. Conversely, hMSCs cultured on tissue culture plastic demonstrated a significant increase in calcium deposition when cultured under osteogenic induction media at all time points.

Similarly, ALP synthesis was significantly greater in hMSCs cultured on POC 1.3-HA composites, relative to hMSCs cultured on PLDLA-TCP control composites and on tissue culture plastic controls under both normal growth media and osteogenic growth media conditions. Interestingly, hMSCs cultured on all experimental and control composite materials in normal and osteogenic growth media demonstrated a significant increase in ALP production relative to cells cultured on control tissue culture plastic (Figure 5d). ^[27,30]^

### 1.17 Transcriptomics profiling of hMSCs cultured on citrate-based composites

To assess the modulation of hMSCs signaling pathways on citrate-based composites, transcriptomics profiling of hMSCs cultured for 7, 14 and 21 days was carried out by stranded transcriptome mRNA sequencing. Transcript counts were used to calculate differentially expressed genes (DEG) for POC 1.1-HA, POC 1.3-HA, and control PLDLA-TCP composites at each time point using DESeq2 relative to hMSCs cultured in normal growth media on tissue culture plastic. Principle component analysis (PCA) of all samples revealed a distinct separation between undifferentiated hMSCs cultured on control tissue culture plastic substrates and hMSCs cultured on experimental and control composite materials (Figure 6a). The number and distribution of differentially expressed genes at each time point is detailed in Figure 6b and 6c respectively.

**Figure 6:**
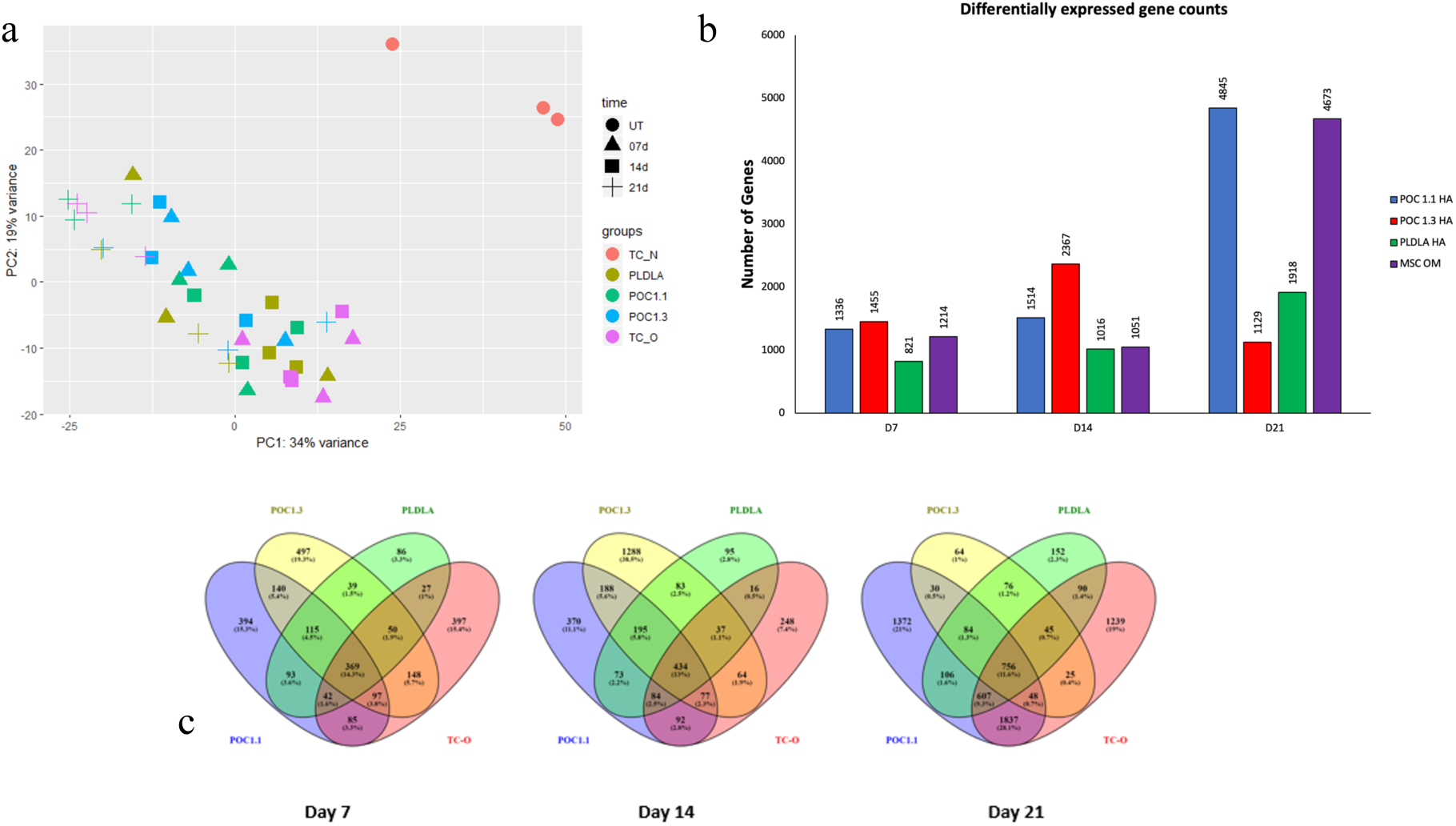
Principle component analysis (PCA) plot for POC 1.1-HA, POC 1.3-HA, and PLDLA-TCP composites showing a clear separation between undifferentiated hMSCs grown in tissue culture plastic, undifferentiated hMSCs grown on citrate-based composites and osteogenic differentiated hMSCs grown in tissue culture plastic (a). The number of differentially expressed genes for each group at each timepoint (b). Venn diagram showing unique and shared differentially expressed genes for all the composites at day 7 day 14 and day 21 (c).

Ingenuity pathway analysis (IPA) was used for the biological interpretation of differentially expressed genes, generating a list of canonical pathways, disease and functions, upstream regulators and mechanistic networks. Pathways with a z score of more than 1.5 were considered as significantly activated while pathways with a z score of less than -1.5 were considered significantly inhibited. Detected enriched gene functions were plotted based on the significance of gene enrichment (-log P-value).

#### 1.17.1 Osteo-specific function analysis

To assess the effects of the investigated composites on osteospecific signaling pathways and bio functions, canonical pathways leading to osteoinduction, osteoconduction, growth factor binding, cellular remodelling and cellular metabolism along with enriched functions related to extracellular matrix, hMSC differentiation, cell attachment, angiogenesis and vasculogenesis were explored for each composite at day 7, 14 and 21 (**Table S1)**.

Critically, the modulation of canonical osteospecific signalling in hMSC populations cultured on POC 1.1-HA, POC 1.3-HA, and control PLDLA-TCP composites at each time point was assessed in detail and activated pathways were classified according to specific osteo-responsive roles, specifically osteoinduction, osteoconduction, growth factor signaling and cellular metabolic pathways (Figure 7a).

**Figure 7:**
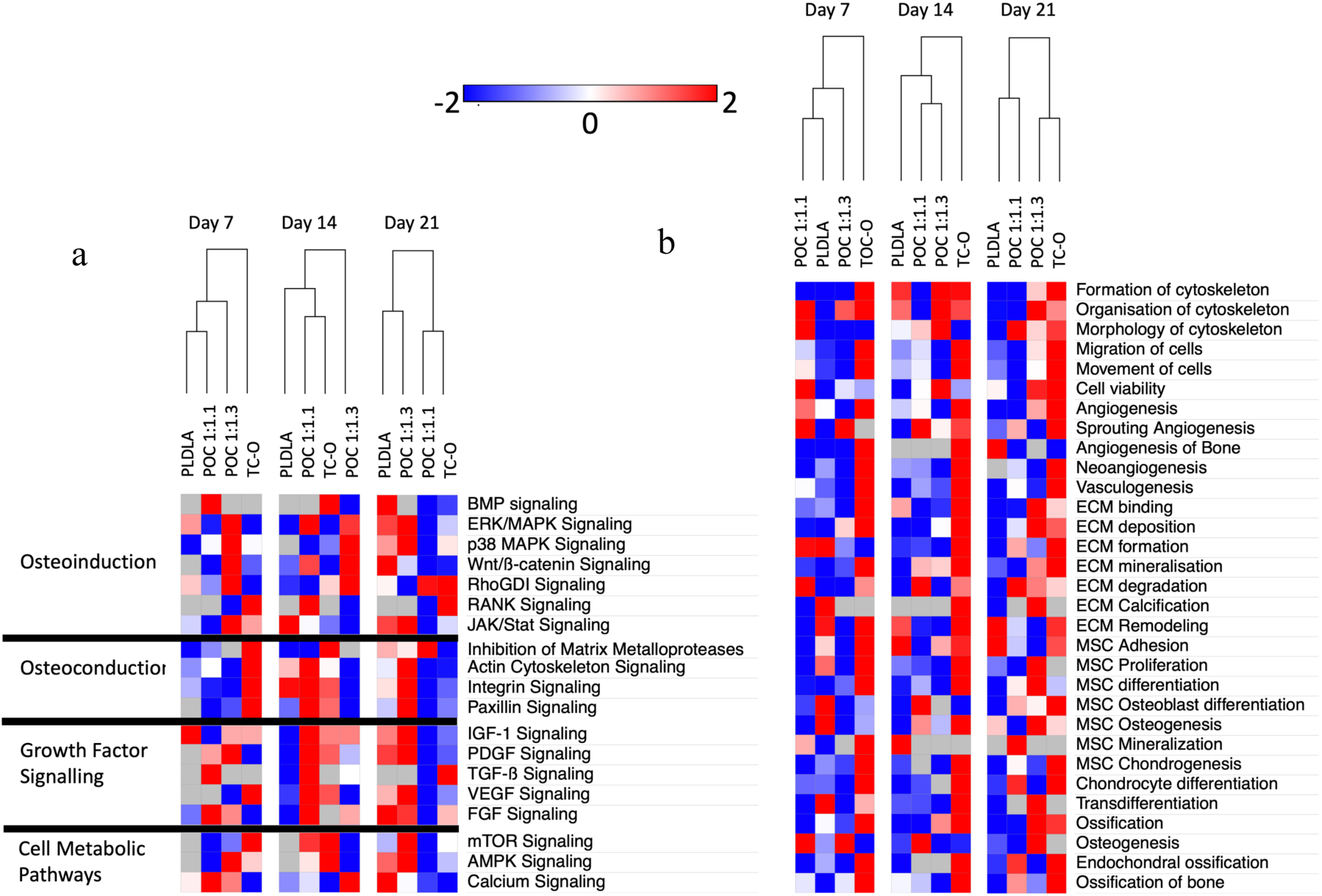
Analysis of the functional response of hMSCs to citrate composite materials. Heat map of osteo-responsive signaling pathways activated by POC 1.1-HA, POC 1.3-HA, and PLDLA-TCP composites (a) and heat map of enriched Bio-functions (b) leading to cytoskeletal rearrangement, cell migration, viability, blood vessel formation, Extracellular matrix related functions, hMSC functions and lineage commitment and ossification related terms activated by citrate-based polymer composites (POC 1:1.1 and POC 1:1.3),and control PLDLA and tissue culture plastic substrates. Plot based on activation score (Z value). A positive Z value represents activation of the biological function, while a negative Z value represents inhibition of the function.

RNA-Seq analysis indicated transcriptional changes in the genes associated with the enrichment of bone morphogenetic protein (BMP), ERK/MAPK, p38 MAPK, RhoGDI, RANK, Wnt/ß-catenin, RhoGDI, and JAK/Stat signaling, pathways attributed to osteoinduction. Specifically, multiple pathways associated with osteoinduction were enriched in MSCs cultured on POC 1:1.3 and control PLDLA composites relative to cells cultured on TCP controls under normal growth conditions.

Osteoconductive pathway enrichment was observed in singnalling processes concerned with cell adhesion and migration such as, Inhibition of MMPs, Actin Cytoskeleton, Integrin, and Paxillin signaling. Similarly, by day 21 the majority of the enrich pathways associated with osteoconductive processes were upregulated in MSCs cultured on POC 1:1.3 and control PLDLA composites relative to cells cultured on TCP controls under normal growth conditions. Although leading to osteoinduction, growth factor signaling pathways were classified separately, and signaling pathways IGF-1, PDGF, TGF-ß, VEGF and FGF signaling pathways were found to be enriched. Again, by day 21 the majority of enriched pathways associated with growth factor signalling were upregulated in MSCs cultured on POC 1:1.3 and control PLDLA composites relative to cells cultured on TCP controls under normal growth conditions.

Metabolic pathways were of interest because the cells undergoing differentiation often require high energy status to fuel the differentiation process which is often marked by increased levels of metabolic activity and activation of metabolic pathways. Enrichment was observed in mTOR, calcium, and AMPK signaling pathways, which were significantly upregulated in MSCs cultured on POC 1:1.3 and control PLDLA composites.

#### 1.17.2 Disease and functions analysis

Bio-functions leading to ECM changes, hMSC lineage commitment to osteoblasts and chondrocytes, ossification, differentiation of chondrocytes and osteoblasts, neoangiogenesis, sprouting angiogenesis, angiogenesis of bone, vasculogenesis, and inflammatory response were explored in MSCs cultured on composite and control substrates for up to 21 days in vitro via stranded transcriptome mRNA sequencing followed by pathway analysis. The number of genes involved in the dataset that corresponds to the above-listed bio functions and their activation state are given in Supplementary Table 1. Enriched bio functions were classified based on their target location and cellular/physiological activity.

The explored enrichment terms were related to ECM formation, binding, mineralization, calcification, and remodeling. hMSC lineage commitment related terms explored were adhesion, adipogenesis, chondrogenesis, osteoblast differentiation, osteogenesis, mineralization, proliferation and differentiation. Enriched bio-functions relating to osteogenesis, endochondral ossification, ossification of bone, neo-angiogenesis, sprouting angiogenesis, and vasculogenesis were also detected (Figure 7b).

A comparison of bio-function activation patterns revealed that hMSCs cultured on tissue culture plastic in osteogenic growth media (TCO) demonstrated the greatest number of activated bio-functions at all explored time points. On day 7, hMSCs cultured on PLDLA-TCP showed the highest activation of ECM calcification, hMSC osteogenesis, and hMSC transdifferentiation. Conversely, hMSCs cultured on POC-HA composites showed the highest activation of cytoskeletal morphology, cell viability, ECM degradation, and osteogenesis. On day 14, hMSCs cultured on the control PLDLA-TCP composite exhibited activation of one bio-function (hMSC adhesion) while hMSCs cultured on POC-HA composites showed the highest activation of 5 (cytoskeletal morphology, sprouting angiogenesis, ECM degradation, osteogenic differentiation of hMSCs, and osteogenesis) and 2 (cell viability and formation of cytoskeleton) bio-functions.

On day 21, hMSCs cultured under TCO conditions showed the highest activation of 19 bio functions. hMSCs cultured on control PLDLA-TCP substrates demonstrated the highest activation of three bio functions (Angiogenesis of bone, MSC adhesion, and ECM remodeling), while hMSCs cultured on POC-HA composites showed the highest activation in 4 bio functions (morphology of cytoskeleton, ECM formation, ECM degradation, and hMSC Mineralisation). hMSCs cultured on POC-HA composites at day 21 showed the highest activation in 8 bio functions (osteogenesis, ossification, Osteogenesis of hMSCs, hMSC proliferation, hMSC differentiation, ECM binding, ECM deposition, and organization of cytoskeleton). TCO hMSCs at 21 days showed the highest activation in 19 bio-functions.

#### 1.17.3 Gene Set Enrichment Analysis (GSEA)

Gene set enrichment analysis was performed using Web-based gene set analysis toolkit (webgestalt), Enriched functions were plotted based on their Normalized Enrichment Score (NES) with 5% FDR cut off. Volcano plots of all Enriched biological processes, cellular component and molecular functions detected for each composite are listed in Supplementary Figure 1. Furthermore, the effects of citrate-based composites on the temporal and spatial activation patterns of Osteospecific canonical pathways signaling pathways in hMSC populations is given in Supplementary Table 2.

### 1.18 In vivo metaphyseal implantation study

#### 1.18.1 Histology and microCT

To assess the biocompatibility and integration of citrate composites in vivo, bone anchor devices were fabricated from the POC 1.1-HA formulation and implanted into the femur and tibia metaphyseal region in a sheep model. A schematic of the bone screw and of the implantation locations can be found in Supplementary Figure 2 and 3 respectively. The ratio of the drilled-generated defect area to the screw area was assessed via sequential uCT imaging of both control PLDLA-TCP anchors and POC 1.1-HA anchors at 3 and 6 months post implantation and of POC 1.1-HA anchors at 12 and 36 months post-implantation (Figure 8a). Two-way ANOVA revealed a significant difference (p=0.025) regarding the effect of time but not the biomaterial formulation on anchor integration. Bone anchor integration was observed to undergo a statistically significant increase at 36-months post-implantation with a ratio of 75.9±7.63% versus the 3-month time point with a ratio of 60.39± 3.74% (p=0.011) (Figure 8b and 8c). Histological analysis of bone tissue indicated that at 12-months post-implantation, the tendon tissue was replaced with trabecular calcified bone tissue at the peri-implant region (Figure 8d).

**Figure 8:**
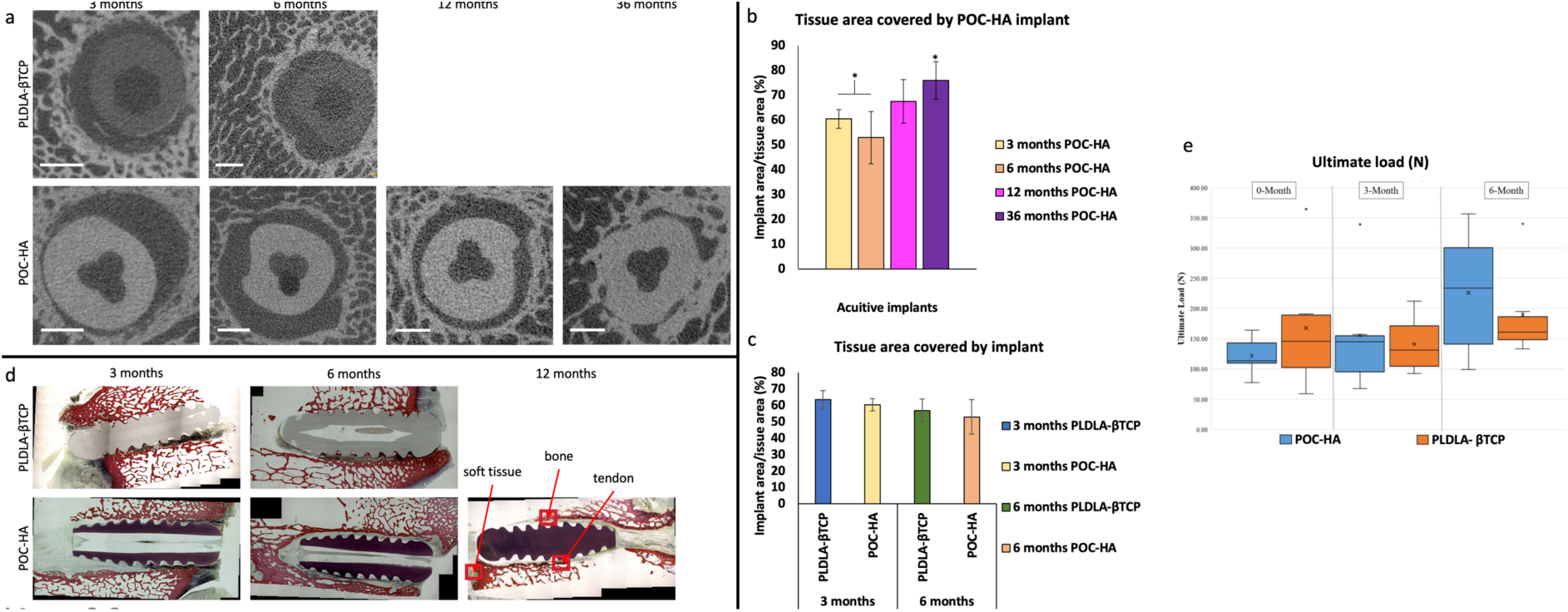
MicroCT and histological analysis of POC and PLDLA-βTCP bone anchor implants. MicroCT images depict the implant inside the tissue cavity, scale bar: 2mm (a). Quantification of peri-implant tissue integration. significant reduction in peri-implant void was observed at 36 months post-implantation of citrate-HA tendon anchor devices (b), however no significant differences were observed between groups at 6 months post-implantation (p<0.05) (c). Histological analysis of the peri-implant region at 12 months post-implantation revealed that the presence of soft tissue was reduced and replaced by bone tissue (d). Ultimate load of Acuitive and Control treatments at the three time points. he “box” is bounded by the first and third quartiles; the “whiskers” represent the maximum/minimum values within the data set; the median data bar, mean data ‘**x**’, and outliers ‘**·**’ are highlighted. There were no significant differences between treatments (p = 0.910), timepoints (p = 0.100), or treatment-timepoint interactions (p = 0.710).

#### 1.18.2 Biomechanical analysis

All biomechanical tests were run to completion. The ultimate load before tendon failure is expressed in Figure 8e. No significant differences in the ultimate load were observed between POC 1:1.1 and PLDAL-TCP groups. The mode of failure (tendon-bone interface, tendon mid-point or tendon-clamp interface) is detailed In supplementary table 3.

## 2 Discussion

Citrate, an intermediator of the tricarboxylic acid cycle, drives the oxidation of multiple nutrients to produce chemical energy in the form of ATP. As well as playing a key role in eukaryote metabolism, the abundance of citrate in bone was first observed by Dickens et al. in 1941, who showed that ∼90% of the body’s citrate is localised within mineralised tissues and makes up ∼1.6% of the bone content. The incorporation of citrate imparts significant biomechanical properties to bone, enhancing the tissue strength and resistance to fracture. Even though the role of citrate in bone apatite crystal stabilization and bone mineralization has been known for several decades, few studies have confirmed the biocompatibility and osteomimetic properties of citrate-derived biomaterials and the ability of citrate to increases cell metabolism in vitro^[31–33]^.

In this study, citrate-based thermoset composites formulated with two different acid to diol ratios (POC 1:1.1 and POC 1:1.3) were synthesized and initially assessed with respect to surface chemistry, degradation rate, and polymer bioceramic interaction. These two citrate-based HA composites were further explored as orthopaedic biomaterials and the ability of these composites to induce osteospecific processes in vitro and bone remodelling in vivo was compared to the thermoplastic composite (PLDLA-TCP), widely used in FDA approved biodegradable orthopaedic devices.

Physicochemical analysis revealed that POC composites underwent polymer hydrolysis at a significantly slower rate to that PLDLA composite materials, which were observed to undergo a non-linear degradation process by DSC analysis. This accelerated degradation of PLDLA composites relative to POC-HA materials was also indicated by an increased surface roughness following a 7 day cell culture study.

Previous studies have reported that citrate-based HA polymer blends and bio-resorbable polymer composites employing a microscale mineral phase enhanced osteoinductivity and osteoconductivity ^[15–18]^, improved surface adhesion of osteoblasts^[21]^, evoked long-term bone and cartilage responses ^[22,23]^ and increased the expression of ALP and osterix^[19]^ in vitro and in vivo. In this study, cytocompatibility analysis of the in vitro response of hMSC populations to POC-HA and control PLDLA-TCP materials indicted that cells cultured on PLDLA-TCP composites showed a statistically significant increase in metabolic activity at 7 and 14 days relative to both POC-HA formulations. Conversely, POC 1.3-HA formulations were observed to induce increased expression of ALP, osteopontin and increased calcium deposition in MSC populations relative to both POC 1.1-HA and control PLDLA-TCP composites.

Transcriptomics profiling by RNA Sequencing has emerged as a powerful tool to elucidate molecular mechanisms underlying biological functions in a cell population. Herein, we profiled transcriptomics changes in hMSCs cultured on POC-HA and control PLDLA-TCP composites using RNAseq and subsequent bioinformatics Ingenuity pathway analysis (IPA). IPA analysis revealed that hMSC population cultured on POC 1.1-HA and POC 1.3-HA composites demonstrated increased activation of multiple signaling pathways and enriched hMSC biological and cellular functions beneficial for osteochondral regeneration and repair.

Transcriptomics analysis of hMSCs cultured on POC-HA composites revealed activation of canonical pathways and enriched molecular/cellular functions relating to osteoinduction and osteoconduction. POC 1.3-HA in particular was found to be the most osteo responsive polymer composite activating 35 (46.6%) of all osteo-responsive pathways detected, while cells alone in osteogenic media, and cultured on POC 1.1-HA and PLDLA-TCP composites positively activated 37.3%, 36% and 28% of all osteo-responsive pathways detected respectively.

Several enriched ontology functions were also detected. These included osteogenesis, ossification, ECM deposition, ECM mineralization and tissue angiogenesis. Interestingly, significant upregulations in pathways associated with the mineralisation of tendon tissues were observed, indicating that citrate materials may be beneficial in the fabrication of tendon anchor devices.

A pilot animal study was conducted to assess the efficacy of a single POC-HA formulation (POC 1:1.1) on tendon anchoring and on peri-implant osteoneogenesis. Due to challenges with manufacturing this was the only POC formulation examined within this study. Although no significant differences is peri-implant bone regeneration were observed between POC and the control PLDLA-TCP group, a significant increase in peri-implant bone formation was observed with the POC anchor following 12 and 36 months in situ. Further biomechanical analysis indicated that again, no significant differences in anchor pull-out forces were observable at 8-weeks post implantation, which agrees with the peri-implant bone deposition data. A major limitation of the animal study however was the lack of a control group post 6 months of implantation, which frustrates the ability to draw definitive conclusions as to the long-term functionality of citrate-HA tendon anchors in vivo. It is encouraging however, that citrate based polymer tendon anchors performed at least as well as control clinically available PLDLA-TCP devices in vivo for up to at least 6 months post-implantation. Further studies will evaluate the in vivo functionality of POC derived anchors fabricated from the more promising 1:1.3 citrate:HA formulation.

In conclusion, this study provides evidence into the osteogenic potential of POC-HA composites and indicates that citrate-based composite materials possess similar or better potential for osteointegration and osteoinduction relative to the industry standard resorbable material PLDLA-TCP.

## 4. Experimental section

### 4.1 Materials

Fully characterized human bone marrow-derived mesenchymal stem cells (hMSCs) from three healthy donors were obtained from collaborating researchers at National University of Ireland, Galway. Bone marrow aspirations were performed at the Translational Research Facility at University Hospital Galway following institutional ethical committee guidelines. Cell culture media and supplements were purchased from Gibco unless stated otherwise. All general chemicals and reagents along with FBS, osteogenic induction media components were purchased from Sigma Aldrich unless stated otherwise. All primary antibodies used in the study were purchased from Abcam. Secondary antibodies and Alamar blue used in the study were purchased from Thermo Fischer. calcein stain was purchased from Invitrogen. Calcium quantification kit was purchased from Stanbio. For ALP quantification, 1-step PNPP substrate was purchased from Sigma Aldrich. For RNA isolation, homogenization columns and isolation kits were purchased from Qiagen. Bioanalyzer chips and reagents were purchased from Agilent. PCR primers were purchased from IDT, Polymerases and other molecular biology reagents were purchased from Promega. Lo bind RNA microcentrifuge tubes were purchased from Eppendorf.

### 4.2 Cell culture

hMSCs were culture expanded in MEM-alpha medium + GlutaMax (Gibco) supplemented with 10% FBS (Sigma Aldrich) and 1% Penicillin streptomycin (Gibco). For osteogenic induction, DMEM low glucose (Sigma Aldrich) supplemented with 100nM dexamethasone, 100Um ascorbic acid 2 phosphate, 10Mm Beta glycerophosphate, 10%FBS and 1% penicillin-streptomycin. All experiments were performed in biological and technical triplicates.

### 4.3 Preparation of citrate-based composites

Citrate-based composites were synthesized as previously reported^[41]^. Briefly, during the pre-polymer synthesis, citric acid (Fisher) and 1,8-octanediol (Sigma) were reacted together in a 1:1.1 acid to diol ratio (POC 1:1.1) or a 1:1.3 acid to diol ratio (POC 1:1.3) at 140C until the desired viscosity was obtained. The uncrosslinked pre-polymer was dissolved in ethanol (Fisher) and composited with hydroxyapatite (Sigma) (60 wt.-%) (POC 1.1-HA or POC 1.3-HA). The composite was mixed until homogeneous, molded into the desired sample shape, and crosslinked at 80C for 7d.

### 4.4 Preparation of PLDLA composites

PLDLA 70/30 (Corbion) was dissolved in dichloromethane (Fisher) and composited with 15 wt.-% tricalcium phosphate (CaP Biomaterials) (PLDLA-TCP). After mixing until homogeneous, the resulting composite was molded into a disk shape.

### 4.5 Compressive mechanical properties

To evaluate the compressive mechanical properties, unconfined compression tests were performed using an Electromechanical Universal testing machine Q Test-150 (MTS, Eden Prairie, MN, USA). Briefly, cylindrical shaped specimens 6×12 mm^2^ (diameter x height) were compressed at a rate of 1.3 mm/min to failure. The initial modulus was calculated by measuring the gradient at 10% of compression of the stress–strain curve. The results werevpresented as mean +/- standard deviation (n = 4).

### 4.6 Composite Degradation

The degradation rate of the 6 x 12 mm^2^ (diameter x thickness) cylindrical samples was assessed in vitro in phosphate buffered saline (PBS) (pH 7.4) at 77C under static conditions. PBS was changed every week to ensure that the pH did not drop below 7. At predetermined time points, samples were extensively rinsed with deionized water and lyophilized before weighing. Weight loss was calculated by comparing the initial weight (W_0_) with the weight measured at predetermined time points (W_t_), as shown in Eq. (1). The results are presented as mean +/- standard deviation (n = 6).

Mass loss (W_0_ - W_t_)/W_0_ * 100% (1)

### 4.7 Analysis of surface topography and roughness

Surface roughness variations were tested using white light interferometry via a Filmetrics Profilm 3D non-contact profilometer (20x objective lens used with a 4x optical zoom). The PLDLA-TCP, POC 1.1 and POC 1.3 substrates were immersed in 1x phosphate buffer saline (PBS) for up to 14 days (D14) and six images were collected for each condition and the average arithmetic mean deviation (Ra), maximum height (Rz) were recorded in the line roughness (1D) mode. To get an overview of the 2D profile the arithmetic mean height (Sa) and root mean square height (Sq) were recorded via area roughness method. Based on these measurements, the surface topography, the roughness variation in both 1D and 2D were analysed.

### 4.8 Cell seeding

hMSCs were seeded on biomaterial samples to assess their viability, metabolic activity, and osteogenic differentiation capacity. Tissue culture plastic was used as a control. The samples were washed in 70% ethanol and soaked overnight in hMSC growth medium in a CO_2_ incubator before cell seeding. hMSCs were seeded at the desired plating density in a volume not exceeding 150 uL. The cell-seeded samples were allowed to stand for 1hr in a CO_2_ incubator at 37^°^C to initiate cell attachment. The samples were then transferred to a 12 well tissue culture plate and carefully supplemented with 3 mL growth medium per well. Growth medium was replaced every three days.

### 4.9 Calcein staining for assessment of cell attachment

hMSCs were seeded at a density of 1×10^5^ cells/cm^2^ per sample in a 12 well tissue culture plate as explained previously. The cells were cultured in complete hMSC growth medium for 7 days before staining for calcein. The working dilution of calcein stain was produced as per the manufacturer’s instructions. The media was aspirated and 750 uL of working dilution of calcein stain was added on top of the composite surface and incubated for 30 min in a CO_2_ incubator at 37^°^C. The cells were subsequently viewed under an inverted fluorescent microscope (Olympus and cell attachment documented in brightfield and FITC channel.

### 4.10 Alamar Blue Metabolic Activity Assay

Metabolic activity of hMSCs was assessed using Alamar blue assay. MSCs were seeded on circular discs of POC (without HA), POC 1.1-HA, POC 1.3-HA, PLDLA-TCP composites, and on tissue culture plastic as described previously. Briefly, 1×10^5^ cells were seeded per sample and cultured for 7, 14, 21 and 28 days in hMSC growth medium to assess the metabolic activity of cells. At the end of each time point, the media was aspirated, the sample was washed in sterile PBS, and 1 ml of Alamar blue reagent (1/10 dilution in complete MSC media) was added to each sample and incubated in a CO_2_ incubator for 5hr at 37^c^. After incubation, the supernatant was collected and transferred to 96 well black plates (200 µl/well) and the fluorescence was measured at 570 nm. Alamar blue reagent added to wells without composite or hMSCs served as the negative control while hMSCs grown in tissue culture plastic (no polymer) served as the positive control. The percentage of metabolic activity was expressed for each composite at the respective time points normalised to cells on tissue culture plastic.

### 4.11 Cell energy phenotype assay; Seahorse analyzer

The OCR and ECAR of hMSCs was measured using the Seahorse Cell Mito Stress Test with the extracellular flux Seahorse XFp analyzer (Agilent), in order to assess the mitochondrial respiration and glycolysis, respectively.^[165, 166]^ Briefly, hMSCs were seeded in PLL-coated Seahorse XF cell culture plates (25 000 cells per well), As per manufacturer’s instructions, 1 h prior to flux analysis, samples were placed at 37 °C in a non-CO_2_ incubator and the media replaced by Seahorse Base XF medium supplemented with 1 mm sodium pyruvate, 2 mm l-glutamine and 10 mm glucose at pH 7.4 for CO_2_ and O_2_equilibration. During this preincubation, the four drugs necessary to perform the Cell Mito Stress Test were loaded into the XF Sensor Cartridges with a preoptimized concentration of 2 µm for the oligomycin, 1 µm of rotenone and antimycin A (provided already mixed by the manufacturer to be loaded and injected together) and 3 µm of carbonyl cyanide-4 (trifluoromethoxy) phenylhydrazone. All the OCR and ECAR values were normalized by DNA content measured for each well using the Quant-iT PicoGreen dsDNA Assay Kit (Invitrogen, Dublin, Ireland) as previously described.^[167]^

### 4.12 Osteogenic Differentiation of MSCs cultured on citrate polymers

hMSC osteogenic differentiation potential was assessed by seeding hMSCs supplemented with either osteogenic induction media or normal hMSC growth media. Briefly, 20,000 cells were seeded on samples in a 12 well tissue culture plate as described above. The cells were cultured for 24 hours on the composite to allow attachment before changing the medium to Osteogenic induction media or Normal MSC growth medium. The cultures were maintained for 7, 14 and 21 days with media replacement every 3 days. Cells cultured on tissue culture plastic served as a control.

Osteogenic differentiation of hMSCs cultured with and without the addition of osteogenic induction media were assessed by staining of calcium deposition by alizarin red and by quantification of calcium deposition and ALP production.

### 4.13 ALP Quantification

ALP activity on cells cultured on citrate-based composites was assessed at 14 days using PNPP assay for ALP quantification. Briefly, the media was aspirated, cells were lysed and scraped off from the polymer using 50ul of 0.2% triton x100 in 100Mm PBS for 10 min. The lysates were transferred to a microcentrifuge tube. In a 96 well plate, 10ul of lysate per well was added. The PNPP substrate was equilibrated to room temperature To each well, 100uL of 1 step PNPP substrate was added and mixed well by gentle agitation. The plate was incubated for 30 min at room temp protected from light. The reaction was arrested by addition of 50 uL, 2N NaOH per well. The absorbance was measured immediately at 405 nm.

### 4.14 Calcium quantification

hMSC calcium deposition was assessed using staining for calcium deposition using alizarin red and by quantifying the deposited calcium. Briefly, the media was removed carefully, and samples were rinsed in PBS thrice. Cells were scraped from the sample surface using 0.5 M HCl and the contents were transferred to a microcentrifuge tube. The solution was incubated overnight at 4^°^C in an incubator shaker, centrifuged to remove any debris, and total calcium was assessed using Stanbio calcium quantification kit following manufacturer’s instructions.

### 4.15 Immunocytochemistry for osteogenic markers

hMSCs were stained for the expression of osteopontin (OPN) and osteocalcin (OCN). Briefly, hMSCs were seeded at a density of 20,000 cells/cm^2^ on each sample using either osteogenic induction medium or normal hMSC medium for 7, 14 and 21 days. Tissue culture plastic was used as a control. Media was replaced every 3-4 days. At the end of each time point, the samples were carefully removed, washed with sterile PBS, and cells were fixed in ice-cold methanol for 20min at -20^0^C. The samples were blocked for 1 hr at room temperature in blocking buffer containing 1% BSA, 22.5mg/ml glycine in PBST (PBS+ 0.5% Tween 20). Primary antibodies for OPN and OCN were added at the desired dilutions. Both osteopontin and osteocalcin antibodies were used at a final concentration of 10µg/ml dilution (1/200 for OCN, 1/100 for OPN) in blocking buffer and incubated overnight at 4^°^C in a humidified chamber. Negative control samples were incubated with blocking buffer instead of the primary antibody. Following overnight incubation, primary antibody was removed carefully, the samples were washed thrice in PBS, secondary antibody was added (1/500 dilution) and incubated at room temperature for 1 hr protected from light.

The samples were fixed on to a glass slide, with the side of cell growth facing away from the glass slide. The side of cell growth side is mounted with a coverslip using fluoroshield with DAPI that stains the cell nucleus blue. The slides were allowed to dry overnight in dark and imaged using a confocal microscope (Olympus Fluoview F1000 system). Images were recorded as z stacks of 1um thickness. Quantification of osteopontin and osteocalcin staining was performed using Image J software.

### 4.16 Transcriptional profiling of MSCs

hMSCs from three healthy donors were used in this study in technical triplicates for each polymer and each time point. A total of 2×10^4^ cells were seeded on each sample and cultured in normal hMSC media for 7, 14 and 21 days. hMSCs cultured in tissue culture plastic that received osteogenic induction media served as the positive control. hMSCs grown in standard hMSC culture media served as the baseline control.

### 4.17 RNA Isolation and integrity check

Total RNA was isolated from hMSCs cultured at day 7, day 14 and day 21 using Qiagen RNAsey plus mini kit following the manufacturer’s instructions. Before loading the lysate on to RNAsey plus columns, the cell lysate was passed through QIAshredder columns to ensure complete homogenization and removal of polymer debris and clumps.

Total RNA was quantified using Agilent RNA nanochips in Bioanalyzer 2100. RNA samples with RIN values >8 were used for library preparation. An exon junction assay was performed using an intron spanning GAPDH primer to generate an amplicon size of 500 bp ensuring the functionality of RNA molecules before library preparation. The integrity and functionality checked RNA samples were shipped to GeneCore, EMBL Heidelberg for library preparation and sequencing in low bind RNA tubes (Eppendorf).

### 4.18 Library preparation and sequencing

Barcoded stranded mRNA-seq libraries were prepared from total RNA (∼200 ng) using the Illumina TruSeq stranded mRNA Sample Preparation v2 Kit (Illumina, San Diego, CA, USA) implemented on the liquid handling robot Beckman FXP2. Obtained libraries that passed the QC step were pooled in equimolar amounts; 1.8 pM solution of this pool was loaded on the Illumina sequencer NextSeq 500 and sequenced uni-directionally, generating ∼500 million reads, each 85 bases long. The resulting fastq files were aligned to human genome v38 and Illumina sequencing adaptors were trimmed using STAR sequence alignment tool.

The resulting aligned .bam files were used to extract UCSC annotated transcript counts using the R (V3.5) Bioconductor (V3.8) package TxDb.Hsapiens.UCSC.hg38.knownGene (V3.4.0). The counts data represents the number of sequence fragments that represented each known gene for each sample.

### 4.19 Differential expression analysis

Once the transcript counts **Table**s were generated, Differential expression analysis was performed from count matrix generated in the previous step using R Bioconductor package DESeq2 (V1.22.2) which implements negative binomial distribution linear models to test differential expression. Differentially expressed genes (DEG) for each composite at each time point was calculated against undifferentiated MSCs (TC-N) with an FDR cut off of 5%. Results were arranged in ascending order of adjusted p-value of significance of fold change (p adj). To stabilize the variations in counts data before DEG calculation, two types of count transformations were implemented in the DESeq2 pipeline, the variance stabilization transformation (VST) and the regularized logarithmic transformation (Rlog). Rlog transformations were chosen for the current study. PCA and MDS plots were used to assess data quality.

### 4.20 Ingenuity pathway analysis (IPA)

IPA (Qiagen) analysis was performed to elucidate canonical pathways, upstream regulator functions along with ontology analysis for diseases and functions. Top canonical pathways and enriched gene functions falling under osteoinduction, osteoconduction, cellular remodelling, growth factor binding, effects on extracellular matrix, angiogenesis, vasculogenesis, cell movement, metabolism and inflammation were identified and explored in detail.

### 4.21 Gene Set Enrichment Analysis (GSEA)

Gene set enrichment analysis was performed to understand the biological processes, cellular component, molecular functions relating to the differentially expressed genes from each group at each time point. Enriched biological functions were analyzed using Web-based gene set analysis toolkit (webgestalt), enriched functions were plotted based on their Normalized Enrichment Score (NES). A false discovery rate (FDR) of 0.05 was used for the analysis. Positive enrichment score represents activation of a function while a negative enrichment score represents inhibition of a function.

### 4.22 In vivo metaphyseal implantation study

For the preclinical study, female skeletally mature (3+ years of age) sheep (ovis aries) of 70-90 kg were used as the animal model due to their similar knee/joint and ACL shape and length anatomy to humans. The animals were treated with control and POC-HA femoral and tibial screws and allocated in timepoints of 3 and 6 months in for a total of 24 animals. Each animal was treated with tibial and femoral interference screws to fix the implanted long digital extensor tendon graft which was selected to replace the animal’s removed ACL ligament. The screws allocated for histological analysis were also scanned with microCT in order to assess bone regeneration. A second animal study was conducted including 6 animals treated only with POC-HA tibial and femoral screws for timepoints of 12 and 24 months with the latter being extended to 36 months after compliance amendments. It has to be mentioned that one animal unexpectedly died at 31 months instead of the 36-month group which it was allocated to. The data are included in the 36-month group for analysis. Sheep knee-joints were excised and underwent microCT analysis and histology processing/histomorphometry imaging.

For the analysis of the microCT images, two open polygon ROIs were selected. One ROI contained the tissue cavity and the second ROI contained the implant as shown in Figure 17. To assess the closure of the wound due to bone formation, a ratio of the two ROIs (implant/tissue cavity) was calculated and expressed as percentage. Lower percentage would denote lower cavity closure thus healing in the anatomical area whereas higher percentage would denote an improved healing process.

For image analysis, 100 images from every ovine patient were analyzed and the average value of their ratio was used for statistical purposes. Due to some timepoints having less than the minimum of N=3 animals, femur and tibia ROI ratio were pooled together in order to have a more robust statistical sample size. The first timepoints of 3 and 6 months that include both control and POC-HA implants were compared with a 2-way ANOVA test to evaluate the impact of time and treatment on bone healing.

Since POC-HA implants were studied until 36 months, a Kruskal-Wallis non-parametric test was used to investigate the effect of time in the wound regeneration, followed by Mann-Whitney U tests to identify statistical significance between different groups of interest (p<0.05). Two donor image groups were excluded from the analysis due to their low quality imaging for identification of ROIs. As a result, for the POC-HA implants, the following number of donors were used: 3 months (N=6), 6 months (N=6), 12 months (N=3), 36 months (N=4)

**Figure 3:**
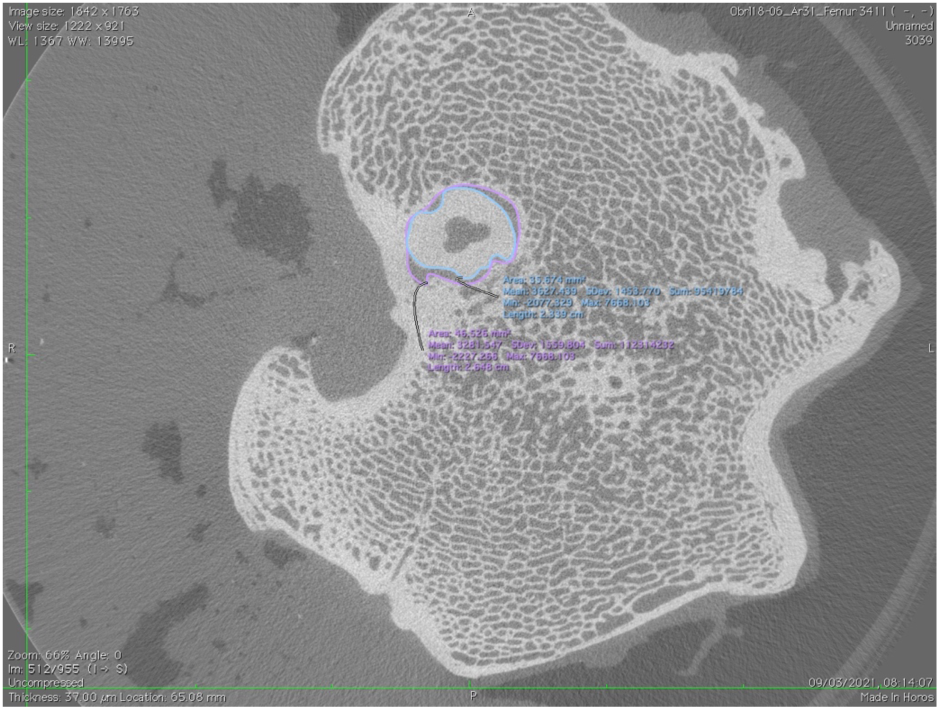
Example of ROIs in a Femur bone

### 4.23 Histological assessment of regeneration

For the histological assessment, bony samples were cut lengthwise down the center of the test article with one half used for decalcified slide preparation and the other half used for undecalcified slide preparation. Both type of tissue samples were fixed in 10% neutral buffered formalin but a different process was selected for each type of sample.

Tissue samples were decalcified with 8% trifluoroacetic acid at room temperature, processed by standard paraffin techniques, cut at 5 μm and stained with Hematoxylin and Eosin (H&E). Undecalcified samples were infiltrated and polymerized into a hardened plastic block (Acrylosin Hard, Dorn and Hart Microedge Inc., Loxley, AL), stained with Sanderson’s Rapid Bone stain to identify differentiated cells and cartilage areas and then counterstained with Van Gieson stain that detects bone and differentiated collagen.

Organ and adjacent soft tissue samples were processed by standard paraffin techniques, cut at 5 μm and stained with H&E. The timepoints used for histology of both control and POC-HA samples at 0 months, 3 months, 6 months and POC-HA only at 12 months.

### 4.24 Destructive biomechanical analysis

Surgically treated knee joints were trimmed at the proximal end of the femur and the distal tibia, leaving the collateral soft tissue ligament across the joint. Samples were kept hydrated via physiologic saline spray at approximately ten-minute intervals during the entire preparation protocol. The collateral ligaments were transected, and the joint capsule opened. The PCL tendon was cut. The ACL tendon was cut as close as possible to the articulating surface of the bone (i.e., the tibia or femur) containing the screw allocated to histology (**Figure 2**). This fine dissection process created two (n = 2) constructs from each sample; one for destructive biomechanics (i.e., the screw-tendon construct embedded in the bone plus the tendon graft length) and one for histology (i.e., the screw-tendon construct embedded in the bone). For samples allocated to destructive biomechanics the bone of interest was potted within a PVC sleeve, using a two-part epoxy resin (Smooth Cast 321, Smooth-on Inc. Easton, Pennsylvania). Prior to potting, several screws were drilled into both the proximal and distal portions of the bone to increase the purchase of the bone within the potting sleeve. The width and thickness of each tendon was measured using a caliper, and cross-sectional area (CSA; mm^2^) of each tendon was calculated by assuming a standard ellipse shape (**Equation 1**). CSA information was utilized to transform force data into the stress space.

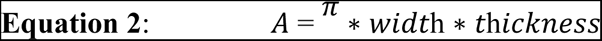

Specimens were mounted into the testing frame (Mini Bionix 858, MTS Systems, Eden Prairie, MN) using specially designed fixtures attached to a 5000 Newton (N) capacity force transducer (Supplementary Figure 4). An upper fixture grip attached the MTS machine’s actuator, was used to clamp onto the ACL tendon. Solid carbon dioxide was laid around the clamp to convert it into a cryo-clamp lowering the temperature to at least -10^ο^C for biomechanical testing. Cryo-clamp temperature was monitored with a thermocouple (Model 52 II, Fluke Corporation, Everett, WA). Specimen hydration was maintained during testing via ambient temperature physiologic saline spray approximately every 10 minutes.

Destructive biomechanical testing included two phases: (A) preconditioning and (B) ramp to failure. Ramp to failure is a destructive test and was performed as the last test in the evaluation sequence. All loads imparted on the samples were applied quasi-statically and aligned collinear to the physiologic loading direction of the tendon. All samples were loaded in the same approximate orientation with respect to bone and tendon orientation.

To minimize the viscoelastic effects on the measured biomechanical response, five (n =5) cyclic tensile loads ranging between 0 and 2% strain were applied for the purpose of preconditioning the tendon. The preconditioning phase was preceded by a ∼2 minute preload phase. A static preload of 10 N was applied to all specimens for ∼2 minutes or until the specimen was fully relaxed. The sample’s reference gauge length was measured as the tendon’s distance, in millimeters, from the bottom of the cryo-clamp’s grip to the tendon’s insertion into the bone (i.e., the tibia or femur) following the 10 N preload. The sample’s reference gauge length was used to transform displacement data into the strain space.

To characterize structural and material properties of the repaired tissue, the specimens were quasi-statically loaded to failure at a rate of 0.5% strain / sec. Load and displacement data was acquired at 100 Hz. Digital videos were collected during ramp to failure testing for failure mode analyses.

## Acknowledgements

This work was funded through Science Foundation Ireland (SFI) and the European Regional Development Fund (Grant No. 13/RC/2073). The authors would like to acknowledge Genecore EMBL, REMEDI University of Galway, TRF University of Galway, Genomics core facility University of Galway, Flow cytometry Core facility University of Galway. The authors also acknowledge the facilities and scientific and technical assistance of the Centre for Microscopy & Imaging at the National University of Ireland Galway.

