## Supplementary Information for "A Functional Analysis of a Resorbable Citrate-based Composite Tendon Anchor"

Supplementary Table 1

To understand the how POCHAs modulate osteo-responsive signaling pathways and bio functions, canonical pathways leading to osteoinduction, osteoconduction, growth factor binding, cellular remodelling and cellular metabolism along with enriched functions related to extracellular matrix, MSC differentiation, cell attachment, angiogenesis and vasculogenesis were explored in detail for each polymer composite at each time point. Details of the enriched osteo-responsive canonical pathways are explained below. A positive z-score indicates a predicted activation, and a negative z-score indicates a predicted inactivation of the enriched pathway. Enriched pathways which do not have an activation score are considered neither activated nor inhibited. List of osteo-responsive pathways and bio functions leading to osteogenic differentiation that were explored in detail are listed in Supplementary Table 1.

| | | **Samples** | | | | | | | | | | | |
|---|---|---|---|---|---|---|---|---|---|---|---|---|---|
| **Pathway function** | **Pathway Name** | **07d_PLDLA** | **07d_POC1.1** | **07d_POC1.3** | **07d_TC_O** | **14d_PLDLA** | **14d_POC1.1** | **14d_POC1.3** | **14d_TC-O** | **21d_POC1.1** | **21d_POC1.3** | **21d_PLDLA** | **21d_TC-O** |
| Osteogenic Canonical | BMP signaling | NA | + | NA | NA | NA | NA | - | NA | - | NA | NA | - |
| | ERK/MAPK Signaling | + | + | + | + | NA | + | + | NA | - | + | + | - |
| | p38 MAPK Signaling | NA | + | + | + | NA | + | + | + | - | + | NA | - |
| | RhoGDI Signaling | - | - | + | - | - | - | + | - | + | - | + | + |
| | RANK Signaling | NA | NA | + | + | NA | + | + | NA | - | NA | NA | - |
| | Wnt/ß-catenin Signaling | NA | - | NA | - | - | + | + | - | - | NA | + | - |
| | Wnt/Ca+ pathway | NA | NA | NA | NA | NA | NA | - | NA | - | NA | - | - |
| | Inhibition of MMPs | + | + | NA | + | + | + | NA | + | + | + | + | + |
| Cellular Remodeling | Actin Cytoskeleton Signaling | - | - | - | + | + | + | - | NA | - | + | - | - |
| | Integrin Signaling | - | - | - | + | + | + | - | + | - | + | - | - |
| | Paxillin Signaling | NA | - | - | + | - | + | - | + | - | + | - | - |

| | | | | | | | | | | | | | |
|---|---|---|---|---|---|---|---|---|---|---|---|---|---|
| Growth factor Signaling | IGF-1 Signaling | + | + | + | + | NA | + | + | + | - | + | + | - |
| | PDGF Signaling | NA | + | + | + | NA | + | + | + | - | + | + | - |
| | TGF-ß Signaling | NA | + | NA | NA | NA | + | + | NA | - | NA | NA | - |
| | VEGF Signaling | NA | NA | + | + | NA | + | - | + | - | + | NA | - |
| | FGF Signaling | + | + | + | + | NA | + | + | NA | - | + | + | NA |
| Cellular Metabolism | mTOR Signaling | NA | - | - | + | NA | + | - | + | - | NA | - | - |
| | Calcium Signaling | + | + | + | + | + | + | + | NA | - | + | + | - |
| | AMPK Signaling | NA | - | + | + | NA | + | - | + | - | + | + | - |
| **Bio-functions** | Angiogenesis | + | - | NA | + | - | - | NA | + | - | NA | NA | - |
| | Vasculogenesis | + | - | NA | - | NA | NA | NA | - | - | NA | NA | - |
| | Activation of bone cells | NA | NA | NA | NA | NA | + | NA | NA | NA | + | NA | NA |
| | Differentiation of osteoblasts | NA | NA | NA | NA | NA | NA | NA | NA | NA | + | NA | NA |
| | Activation of osteoclasts | + | - | NA | NA | NA | + | NA | NA | NA | + | + | NA |
| | Organization of cytoskeleton | NA | - | - | + | + | + | - | + | - | + | - | - |
| | Formation of cytoskeleton | NA | NA | NA | + | + | NA | - | + | - | NA | NA | - |

*Table 1: List of osteo-responsible canonical pathways and bio functions modulated by CBPBHAs. Pathways and bio-functions with positive z score are represented by +, with negative z score are represented by – and ones that were not present in the dataset are represented as NA.*

Supplementary Figure 1. Gene set enrichment analysis was performed using Web-based gene set analysis toolkit (webgestalt), Enriched functions were plotted based on their Normalized Enrichment Score (NES) with 5% FDR cut off. Volcano plots of all Enriched biological processes, cellular component and molecular functions detected for each polymer composite are listed below.

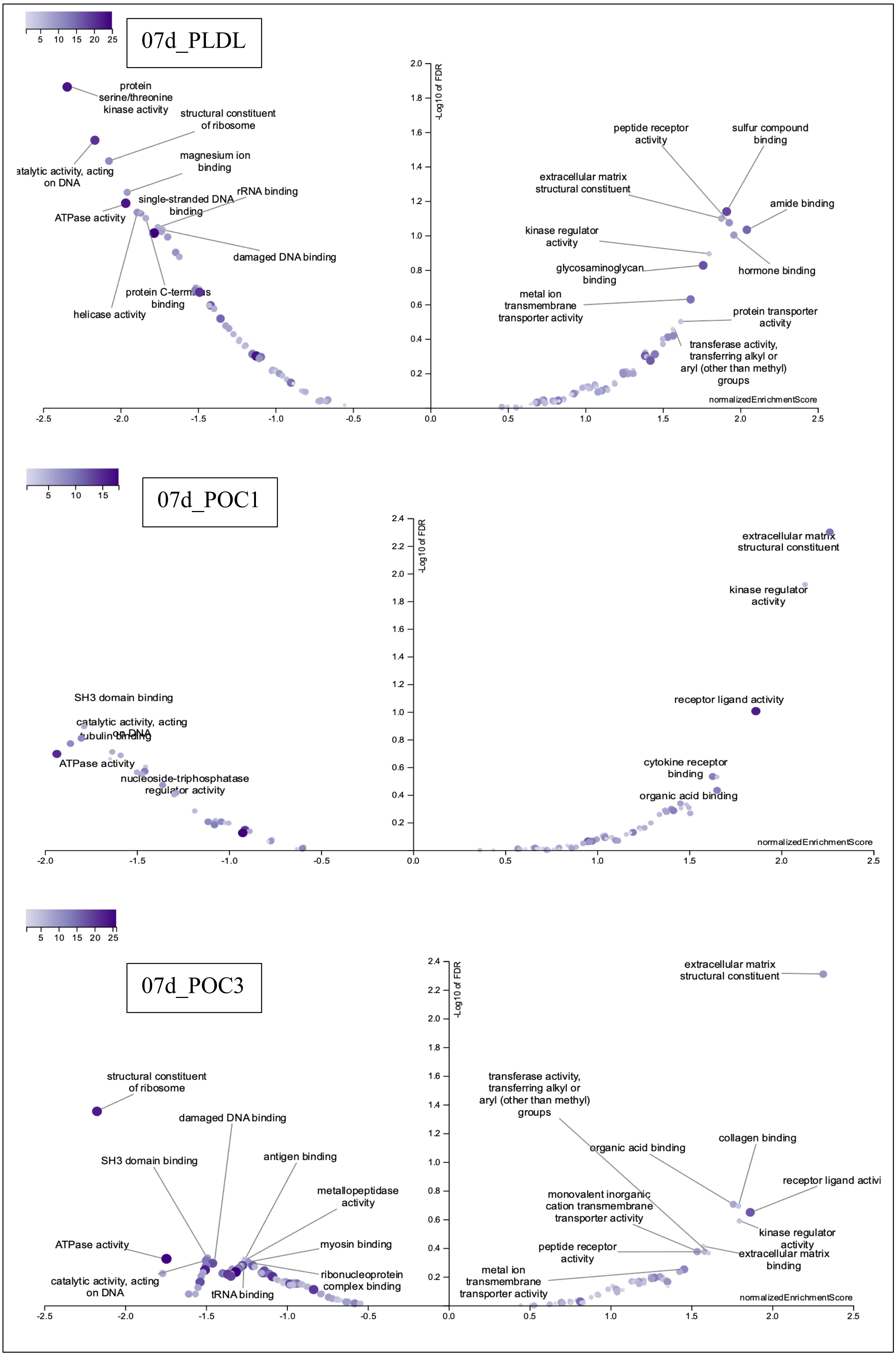

07d_PLDL
-Log10 of FDR
normalizedEnrichmentScore
protein serine/threonine kinase activity
structural constituent of ribosome
atalytic activity, acting on DNA
magnesium ion binding
single-stranded DNA binding
rRNA binding
ATPase activity
damaged DNA binding
helicase activity
binding
peptide receptor activity
sulfur compound binding
extracellular matrix structural constituent
amide binding
kinase regulator activity
hormone binding
glycosaminoglycan binding
metal ion transmembrane transporter activity
protein transporter activity
transferase activity, transferring alkyl or aryl (other than methyl) groups
07d_POC1
-Log10 of FDR
normalizedEnrichmentScore
extracellular matrix structural constituent
kinase regulator activity
SH3 domain binding
catalytic activity, acting
ATPase activity
nucleoside-triphosphatase
receptor ligand activity
cytokine receptor binding
organic acid binding
07d_POC3
-Log10 of FDR
normalizedEnrichmentScore
extracellular matrix structural constituent
structural constituent of ribosome
damaged DNA binding
SH3 domain binding
antigen binding
metallopeptidase activity
ATPase activity
myosin binding
catalytic activity, acting on DNA
ribonucleoprotein complex binding
tRNA binding
transferase activity, transferring alkyl or aryl (other than methyl) groups
organic acid binding
collagen binding
receptor ligand activi
monovalent inorganic cation transmembrane transporter activity
kinase regulator activity
peptide receptor activity
extracellular matrix binding
metal ion transmembrane transporter activity

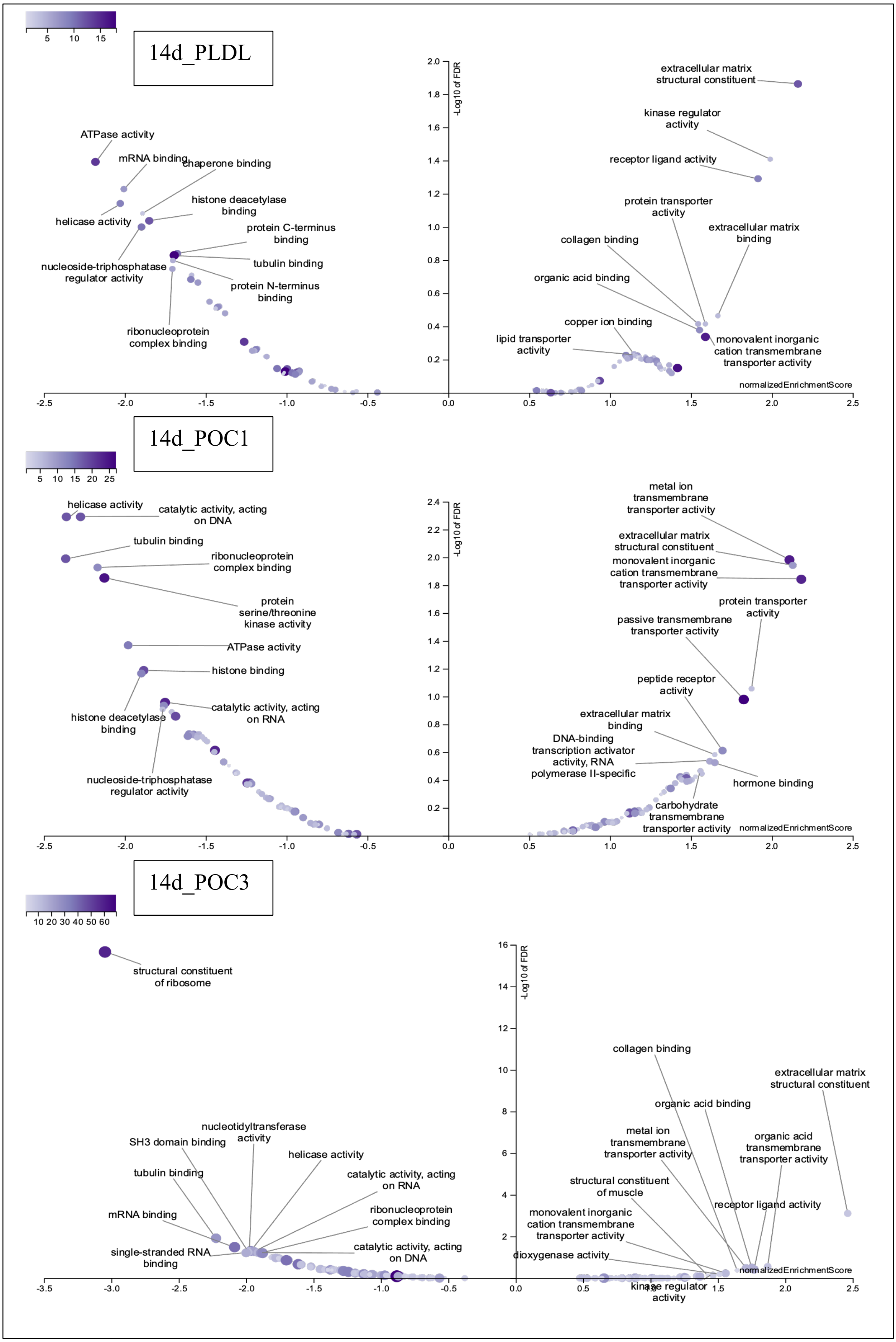

14d_PLDL
extracellular matrix structural constituent
kinase regulator activity
receptor ligand activity
protein transporter activity
extracellular matrix binding
collagen binding
organic acid binding
copper ion binding
lipid transporter activity
monovalent inorganic cation transmembrane transporter activity
ATPase activity
mRNA binding
chaperone binding
helicase activity
histone deacetylase binding
protein C-terminus binding
tubulin binding
nucleoside-triphosphatase regulator activity
protein N-terminus binding
ribonucleoprotein complex binding
-Log10 of FDR
normalizedEnrichmentScore
14d_POC1
helicase activity
catalytic activity, acting on DNA
tubulin binding
ribonucleoprotein complex binding
protein serine/threonine kinase activity
ATPase activity
histone binding
histone deacetylase binding
catalytic activity, acting on RNA
nucleoside-triphosphatase regulator activity
metal ion transmembrane transporter activity
extracellular matrix structural constituent
monovalent inorganic cation transmembrane transporter activity
protein transporter activity
passive transmembrane transporter activity
peptide receptor activity
extracellular matrix binding
DNA-binding transcription activator activity, RNA polymerase II-specific
hormone binding
carbohydrate transmembrane transporter activity
14d_POC3
structural constituent of ribosome
nucleotidyltransferase activity
SH3 domain binding
helicase activity
tubulin binding
catalytic activity, acting on RNA
ribonucleoprotein complex binding
mRNA binding
single-stranded RNA binding
catalytic activity, acting on DNA
collagen binding
extracellular matrix structural constituent
organic acid binding
metal ion transmembrane transporter activity
organic acid transmembrane transporter activity
structural constituent of muscle
receptor ligand activity
monovalent inorganic cation transmembrane transporter activity
dioxygenase activity
kinase regulator activity

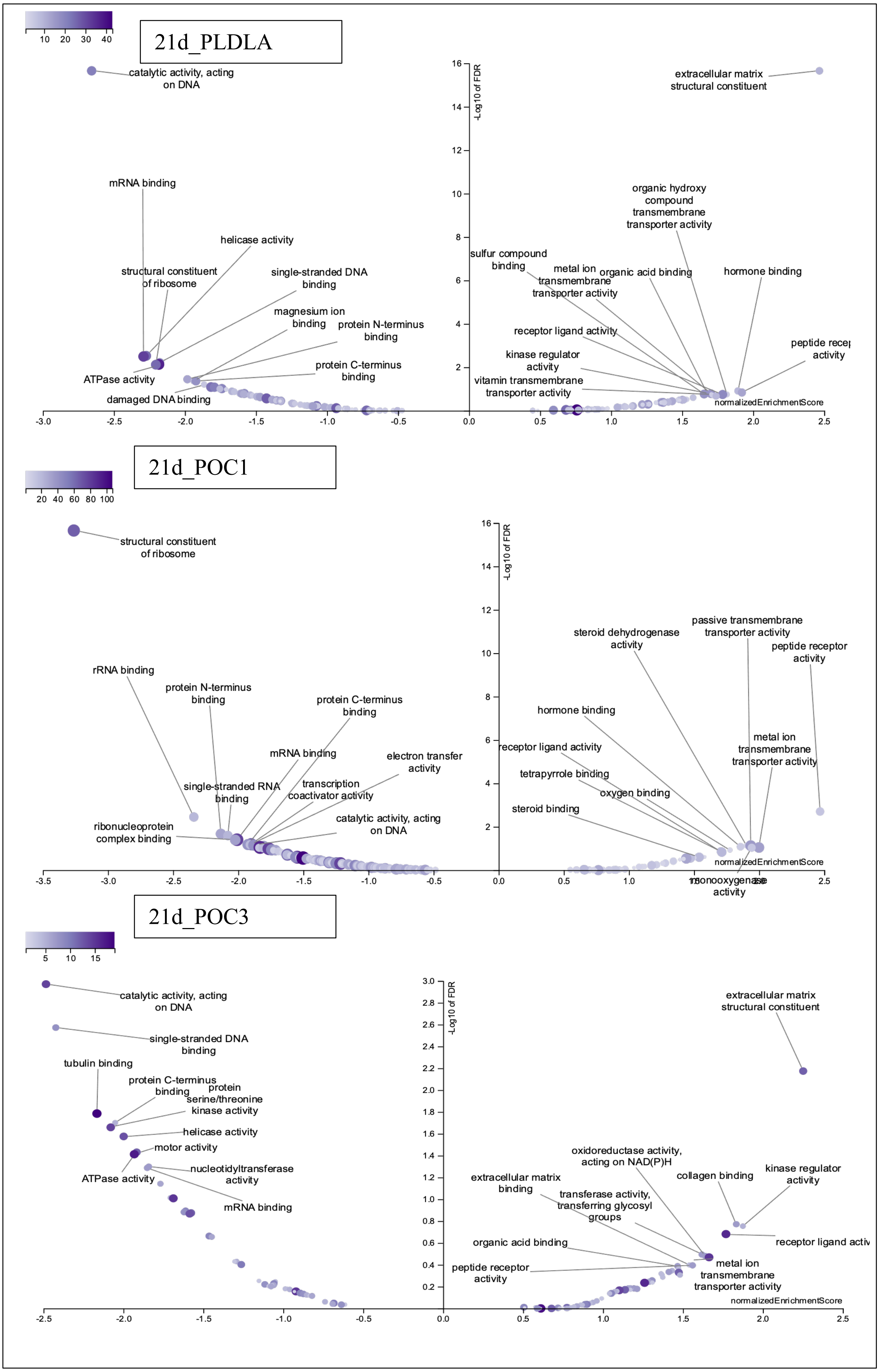
21d_PLDLA
10 20 30 40
catalytic activity, acting on DNA
extracellular matrix structural constituent
-Log10 of FDR
mRNA binding
helicase activity
structural constituent of ribosome
single-stranded DNA binding
magnesium ion binding
protein N-terminus binding
protein C-terminus binding
ATPase activity
damaged DNA binding
organic hydroxy compound transmembrane transporter activity
sulfur compound binding
metal ion transmembrane transporter activity
organic acid binding
hormone binding
receptor ligand activity
kinase regulator activity
vitamin transmembrane transporter activity
peptide recep activity
normalizedEnrichmentScore
21d_POC1
20 40 60 80 100
structural constituent of ribosome
rRNA binding
protein N-terminus binding
protein C-terminus binding
mRNA binding
electron transfer activity
single-stranded RNA binding
transcription coactivator activity
ribonucleoprotein complex binding
catalytic activity, acting on DNA
steroid dehydrogenase activity
passive transmembrane transporter activity
peptide receptor activity
hormone binding
receptor ligand activity
metal ion transmembrane transporter activity
tetrapyrrole binding
oxygen binding
steroid binding
monooxygenase activity
21d_POC3
5 10 15
catalytic activity, acting on DNA
single-stranded DNA binding
tubulin binding
protein C-terminus binding
protein serine/threonine kinase activity
helicase activity
motor activity
ATPase activity
nucleotidyltransferase activity
mRNA binding
extracellular matrix structural constituent
oxidoreductase activity, acting on NAD(P)H
extracellular matrix binding
collagen binding
kinase regulator activity
transferase activity, transferring glycosyl groups
receptor ligand activ
organic acid binding
peptide receptor activity
metal ion transmembrane transporter activity

Supplementary Table 2. Total number of Osteospecific canonical pathways activated by each composite at each time point are as follows.

| | PLDLA | POCHA1:1.1 | POCHA1:1.3 | TCP |
|---|---|---|---|---|
| **Day 7** | **7** | **8** | **10** | **16** |
| **Day 14** | **5** | **18** | **10** | **11** |
| **Day 21** | **9** | **1** | **15** | **1** |
| **Total** | **21** | **27** | **35** | **28** |
| **Percentage** | **28** | **36** | **46.66667** | **37.33333** |

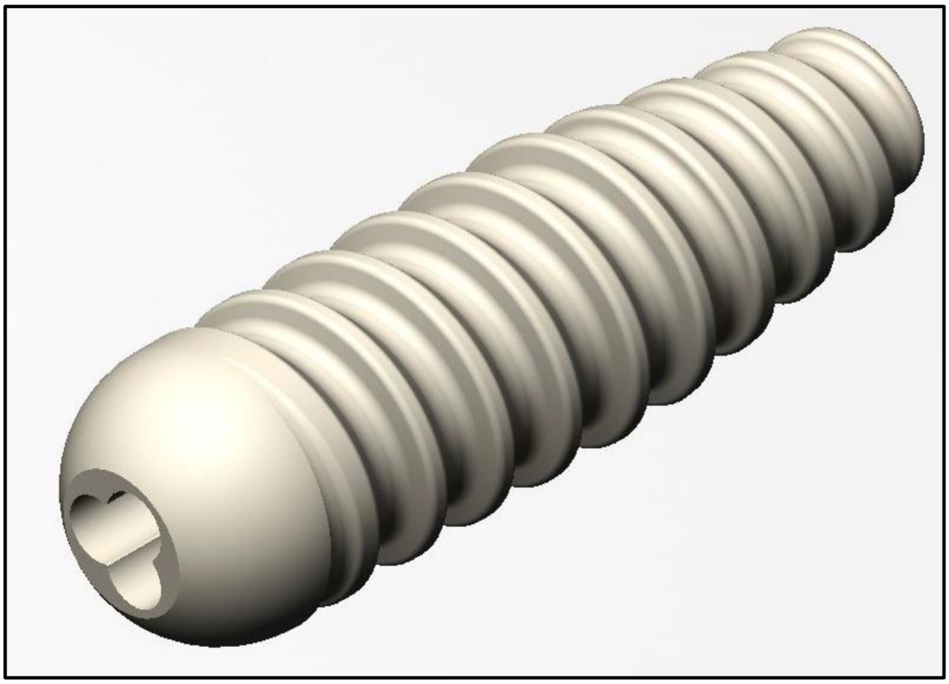

Supplementary Figure 2. Schematic of the tendon anchor screw

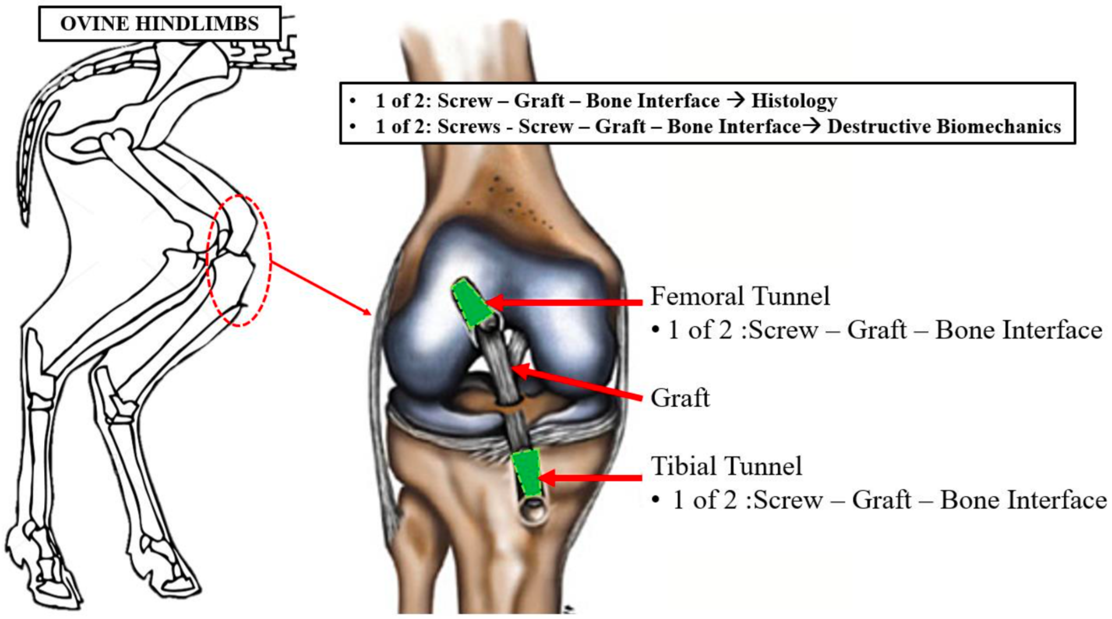


Supplementary Figure 3. Schematic of sheep hind quatre detailing screw locations, and post sacrifice sample allocation.

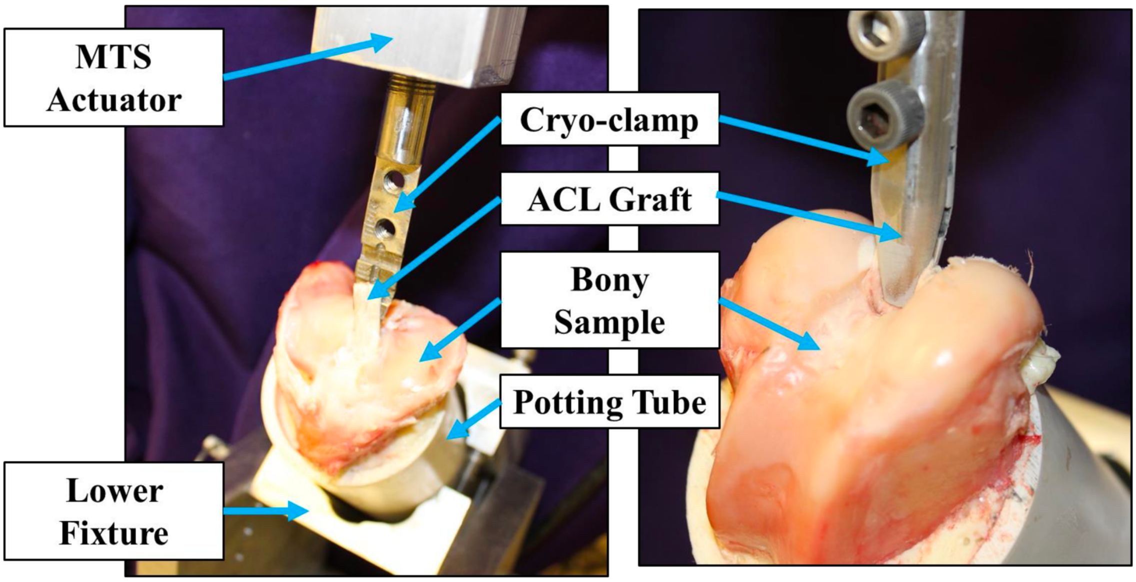


Supplementary Figure 4. Digital image showing the mechanical testing fixture. Components of interest are indicated

The mode of failure is detailed In supplementary table 3. MOF was scored as:

- Tendon-grip interface failure (i.e., yielding / failure of the tendon at its attachment to the mechanical clamp).

- Tendon mid-substance failure (i.e., yielding / failure of the tendon in the center region away from its boney insertion or attachment to the mechanical clamp).

- Tendon-bone interface failure (i.e., yielding / failure of the tendon at its attachment to the bone not necessarily including slippage of the tendon from the bone tunnel).

Supplementary Table 3. Biomechanics testing mode of failure

| | **Tendon Mid-Substance** | | **Tendon-Bone Interface** | | **Tendon-Grip Interface** | |
|---|---|---|---|---|---|---|
| | Number of Specimens | Percent of Specimens (%) | Number of Specimens | Percent of Specimens (%) | Number of Specimens | Percent of Specimens (%) |
| 0 month POC 1:1.1 | 1 | 17% | 5 | 83% | 0 | 0% |
| 0 month PLDLA | 3 | 50% | 2 | 33% | 1 | 17% |
| 3 month POC 1:1.1 | 1 | 17% | 5 | 83% | 0 | 0% |
| 3 month PLDLA | 4 | 67% | 0 | 0% | 2 | 33% |
| 6 month POC 1:1.1 | 1 | 17% | 5 | 83% | 0 | 0% |
| 6 month PLDLA | 2 | 33% | 4 | 67% | 0 | 0% |
